# Machine Learning Optimization of Photosynthetic Microbe Cultivation and Recombinant Protein Production

**DOI:** 10.1101/2021.08.06.453272

**Authors:** Caitlin Gamble, Drew Bryant, Damian Carrieri, Eli Bixby, Jason Dang, Jacob Marshall, David Doughty, Lucy Colwell, Marc Berndl, James Roberts, Michael Frumkin

## Abstract

**Background:** *Arthrospira platensis* (commonly known as spirulina) is a promising new platform for low-cost manufacturing of biopharmaceuticals. However, full realization of the platform’s potential will depend on achieving both high growth rates of spirulina and high expression of therapeutic proteins.

**Objective:** We aimed to optimize culture conditions for the spirulina-based production of therapeutic proteins.

**Methods:** We used a machine learning approach called Bayesian black-box optimization to iteratively guide experiments in 96 photobioreactors that explored the relationship between production outcomes and 17 environmental variables such as pH, temperature, and light intensity.

**Results:** Over 16 rounds of experiments, we identified key variable adjustments that approximately doubled spirulina-based production of heterologous proteins, improving volumetric productivity between 70% to 100% in multiple bioreactor setting configurations.

**Conclusion:** An adaptive, machine learning-based approach to optimize heterologous protein production can improve outcomes based on complex, multivariate experiments, identifying beneficial variable combinations and adjustments that might not otherwise be discoverable within high-dimensional data.

## INTRODUCTION

Biologic drugs have been transforming patient lives for decades. But the traditional systems used to produce therapeutic proteins require complex and costly manufacturing. As a result, affordable and widespread access to biologics remains a challenge^1–3^.

The photosynthetic cyanobacterium *Arthrospira platensis* (commonly known as spirulina) has many advantages compared with traditional platforms used to produce biopharmaceuticals. These include simple, inexpensive growth and downstream processing; photosynthetic metabolism; and Generally Regarded as Safe (GRAS) status with the FDA^4–6^. Genetic engineering of spirulina allows for stable expression of heterologous proteins, including diverse anti-pathogen proteins such as active vaccine antigens, antibodies, and therapeutic enzymes^7^. Together, these features offer the potential to produce disruptively low-cost biologics and biologic cocktails.

A key aim of any emerging biotechnology platform is the production of recombinant proteins at high yields^8^. Environmental variables play a crucial role in achieving high yields of both biomass and heterologous protein^3^. For platforms like spirulina, many continuous environmental variables can be adjusted simultaneously. These include light intensity and spectrum, mixing, aeration, and temperature. Methods to predict the optimal combination of so many variables are desirable.

Historically, the biotechnology industry used one-variable-at-a-time (OVAT) experiments to optimize production medium^9^. But this method is slow, expensive, and laborious, leading to widespread adoption of ‘multivariate data analysis’ (MVA) to improve manufacturing of biologics^10–12^. MVA carried out with a ‘design of experiments’ (DOE) approach has been widely used by biologics manufacturers to optimize production media and growth conditions^10–13^. One biotechnology manufacturer used iterative statistical modeling to increase productivity 34% in mammalian cell lines^14^. However, a central challenge remains in selecting variable subsets and the appropriate variable levels to test, given limited *a priori* knowledge and resources.

Machine learning (ML) approaches can facilitate the adaptive exploration of complex multivariate spaces and can exploit complex, nonlinear, higher-order relationships^15,16^. Among such ML approaches, batched Bayesian optimization (BO) has become a preferred black box method to tune high-complexity systems when trials are expensive to run, observations are noisy, or derivatives of the objective to optimize are unavailable^17,18^. It is a powerful tool to adaptively select experiments and approach an optimum as quickly as possible - even in non-convex multimodal search spaces - while being robust with respect to poorly chosen variable bounds, which may be expensive or impossible to know with accuracy before experimentation^18^.

The ability to build genetically modified spirulina cell lines while tightly controlling photobioreactor environments presents a unique opportunity to adaptively optimize conditions for low-cost production of therapeutics. We report below on the development of a high-throughput experimentation pipeline using custom bioreactor arrays. We describe how an ML-based approach iteratively guided a series of experiments to search across possible combinations of 17 environment variables, which ultimately identified several key variable combinations to improve production outcomes. Finally, we describe how we applied these conditions to cultures of therapeutic strains in larger-scale bioreactors to improve therapeutic protein yield.

## METHODS

See Supplemental Materials.

## RESULTS

### Custom bioreactor arrays allow for high throughput experimentation

We set out to improve volumetric productivity (heterologous protein yield per unit of culture volume and per unit of time) in the spirulina expression system. To enable rapid quantification of heterologous protein for ML-guided optimization, we engineered a spirulina strain (SP699) constitutively expressing green fluorescent protein (GFP) with a C-terminus dimerization scaffold and polyhistidine-tag. This design was similar to our fusion protein designs for therapeutic antibodies. Once fully segregated to homozygosity, cells in the engineered spirulina filaments showed diffuse and uniform GFP fluorescence (**Figure 1A**). Using western blot, we found that the GFP fusion protein comprised 2.7% of total protein (**Supplemental Figure S1**). This level was within the typical range for antibodies in development as therapeutics^7^.

**Figure 1:**
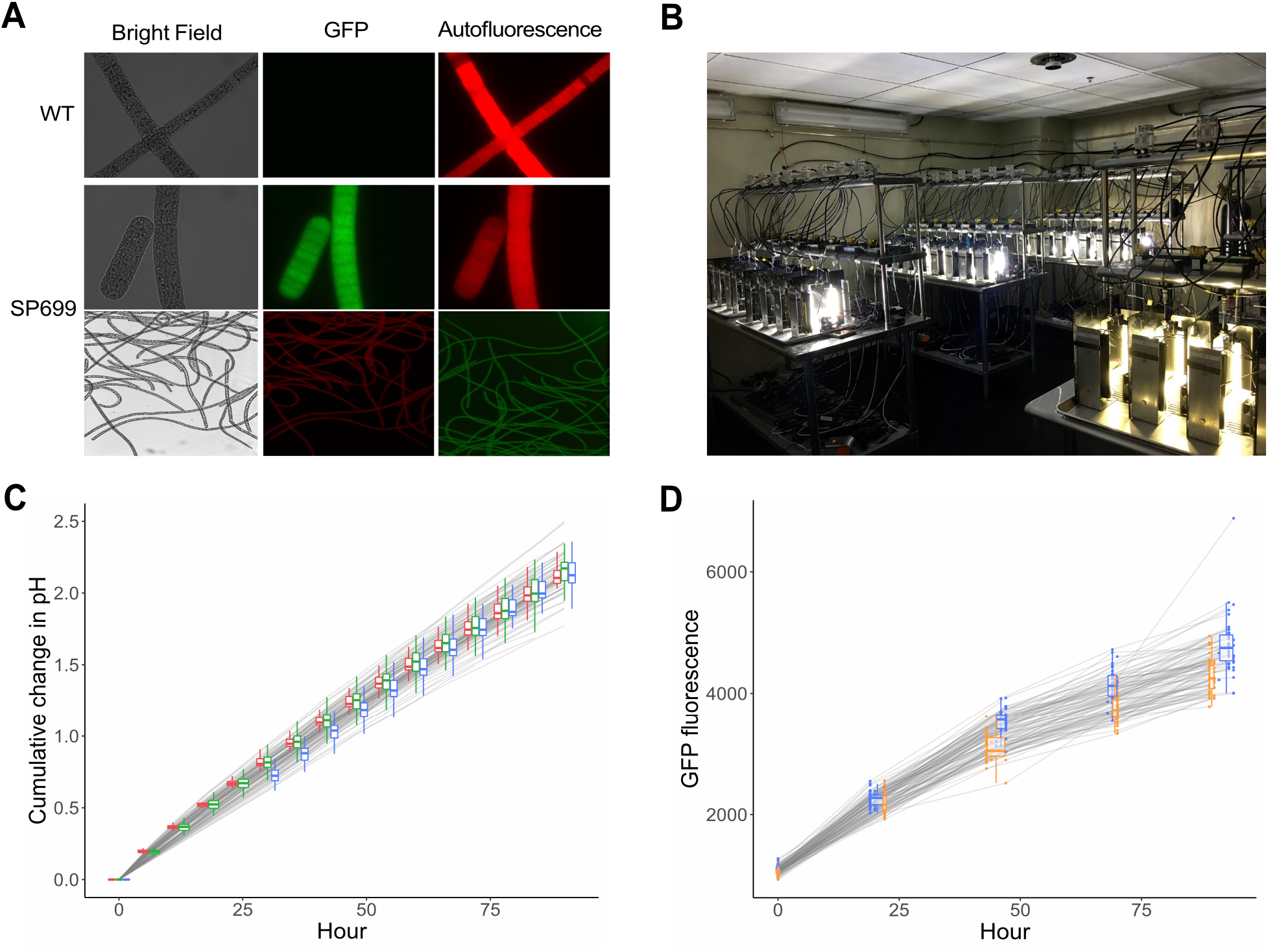
Bioreactor facility setup and commissioning establishes foundation for high-throughput exploration of culturing conditions. **A)** Microscope images of WT (UTEX LB 1926) and GFP-scaffold (SP699) spirulina strains shown in bright field, with Texas Red filter, and with YFP filter shown at 1000x magnification (top) and 100x magnification (bottom) using a Leica DM5000B microscope. **B)** Photobioreactors in a facility of 96 independently controlled reactors. Each reactor is independently controlled with programmed settings. **C and D)** Bioreactor commissioning runs as time series of cumulative change in pH (left plot, total n = 61), and GFP fluorescence (right plot, total n = 93). Box plots depict median and interquartile range by run set (color) and timepoint. Precise sampling times for GFP readings vary. Points reflect actual sample times; box plots are grouped by day.

To carry out high-throughput experimentation, we constructed 96 bioreactors each with a liquid capacity of 0.5 liters. This reactor array was designed with a light path-length similar to commercial-sized reactors to facilitate scale-up of improved growth regimens. Each bioreactor was constructed with independently controlled, continuously tunable settings for LED-illumination, temperature, airflow (for mixing), and CO_2_ flow (for pH control). Initial growth conditions were selected to closely mimic commercial and laboratory growth conditions for *Arthrospira*. The growth media used is based on the ATCC recommended medium 1679 for spirulina: SOT medium for spirulina, modified by doubling nitrate concentration from 2.5 g/L to 5.0 g/L to support higher total biomass accumulation without reaching nitrogen limitation. A growth temperature of 35°C and cultivation pH of 9.75 to 9.95 were selected to closely match published commercial spirulina growth conditions^19^. Cultures were illuminated with constant light at 1000 µmol/m^2^/s from both sides of the bioreactor to mimic existing capabilities of our production-scale system for therapeutics. We defined these initial growth conditions as our “standard” for the study. We next sought to commission the bioreactors by verifying the reproducibility of run outcomes. Starting with inoculation densities of 0.5 g/L, we carried out initial run sets of 15 to 48 bioreactor cultures with the GFP fusion strain. These cultures were grown for ∼90 hours in our standard condition. Throughout each run, we monitored changes in pH and used the cumulative change as a proxy metric for total fixed carbon and, hence, biomass growth (see **Methods**, “**Cumulative pH calculation**”). For each day of the run set, we also sampled the cultures and measured both GFP fluorescence and cell autofluorescence per unit of culture volume (see **Supplemental Figure S2** for strain spectral analysis).

We initially observed a high degree of variance in bioreactor production run metrics. Cumulative pH readings for an early run set of 22 bioreactors had a relative standard deviation of ∼16% at 72 hours. However, with improved processes, especially more uniform airflow rates and tighter control of CO_2_ delivery, we reduced overall variance to a relative standard deviation of ∼6% in cumulative pH change across 61 reactor runs in 3 run sets (**Figure 1C**). GFP fluorescence readings initially had a relative standard deviation of ∼13% at ∼72 hours. With fluorescence normalizations (see **Methods**), we reduced the noise and achieved a relative standard deviation of ∼5-6% within single run sets and ∼8% across 93 reactor runs in 2 run sets (**Figure 1D**). Based on the consistency of these results, we initiated experiments with varying conditions.

In our first set of bioreactor experiments, we varied the intensity of light, a key variable in photosynthetic growth. If carbon fixation rates are less than the cell’s photosynthetic capacity, then increasing the intensity of light can increase growth rates. However, delivery of light amounts that exceed the cell’s capacity to use it for photosynthesis confers no additional growth benefit and can lead to cell damage through the generation of reactive oxygen species. We carried out triplicate bioreactor runs at 13 constant light settings, ranging from ∼120 to 2415 µmol/m^2^/s. We found that cultures grown at higher light intensities had consistently better growth and protein production up to a light intensity around 2000 µmol/m^2^/s. Cultures grown at light intensities above our standard condition of 1000 µmol/m^2^/s had an average of 23% to 38% higher GFP yield at the final time point (group mean = 34% more signal, p-value = 1.8e-05) (**Figure 2A**). Volumetric productivity differences with fluxes of 680 to 1000 µmol/m^2^/s emerged around the 48-hour sampling timepoint. Plotting these results as high-resolution light response curves revealed a plateau of carbon fixation rates per unit volume at about 2000 µmol/m^2^/s, while GFP accumulation rates per unit volume plateaued at a lower intensity of around 1250 µmol/m^2^/s (**Figure 2B**). The results suggested that increases in light intensity between ∼1250 and 2415 µmol/m^2^/s reduces the concentration of heterologous protein per unit of biomass (“potency”), since the rate of carbon fixation per unit of volume continued to increase without similar increases in the rate of heterologous protein accumulation per unit of volume. These results further suggested that our cultures reached their maximum photosynthetic capacity at a light intensity below the maximum achievable with our light equipment. Thus, further improvements in productivity would need to come from other variables.

**Figure 2:**
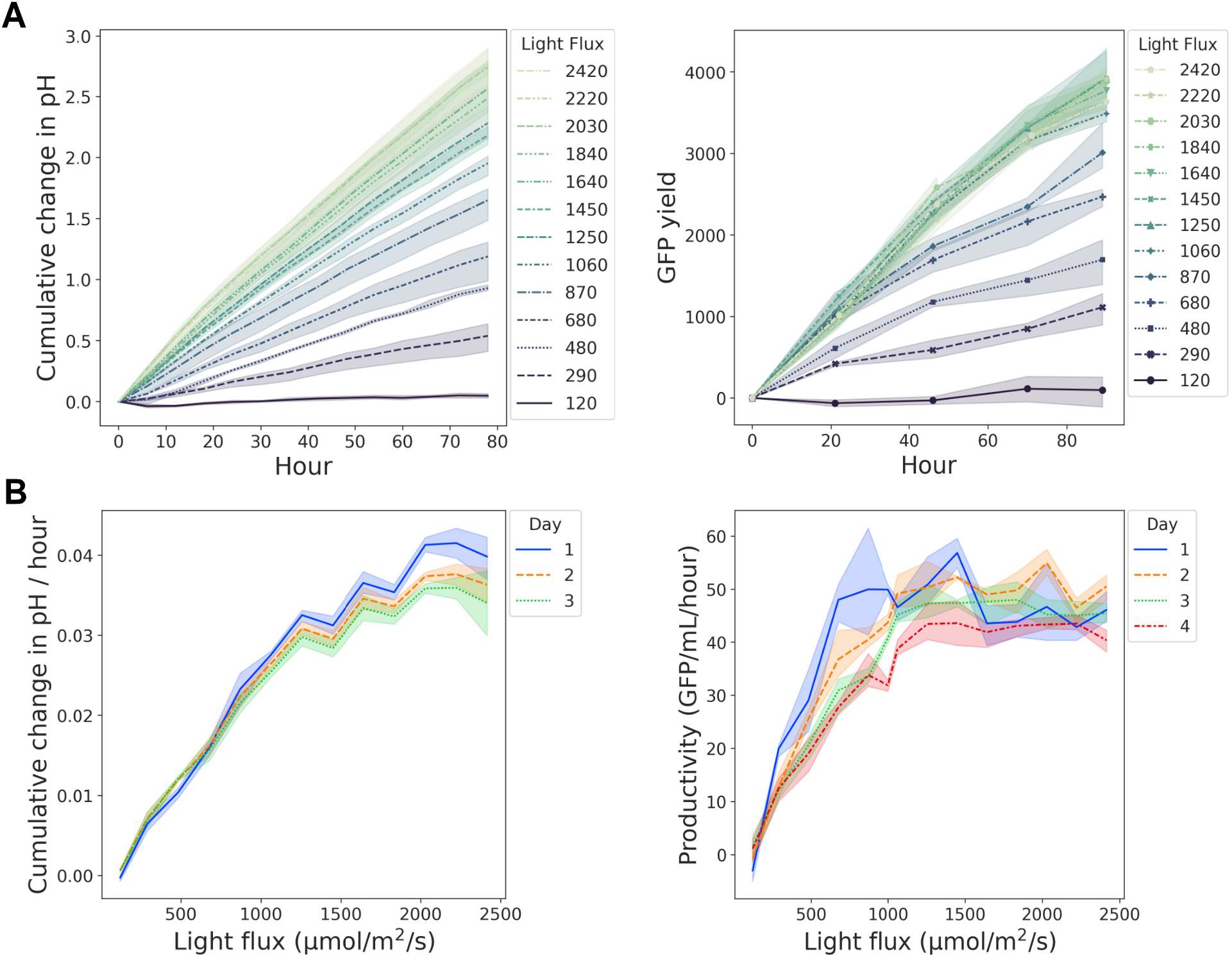
Higher light intensities lead to more culture growth and GFP production until the relationships plateau. **A)** Time series of mean cumulative change in pH (left plot) and mean GFP yield in fluorescence units (right plot) for bioreactor cultures grown at constant light across 13 different target light fluxes (120 - 2415 µmol/m^2^/s). Shaded regions represent the 95% confidence interval based on triplicate runs. **B)** Light response curves for the rate of cumulative pH change (left) and volumetric GFP productivity (right) on each day of the run set (colors). Shaded regions represent the 95% confidence interval based on triplicate runs.

### Bayesian black box mapping of parameter relationships leads to efficient selection of culture conditions with improved performance

As a photosynthetic microbe, spirulina is highly attuned to light characteristics such as intensity, color, and cycling^20^. Thus, we sought to vary these light characteristics in our search for more optimal conditions. We defined 6 different light parameters. Configuration of these parameters specified 1-2 light intensity levels for two different LED types: a set of red-shifted white LEDs (3000 - 4750 Kelvin) and a set of blue-shifted white LEDs (4750 - 6500 Kelvin). For red-shifted LEDs, the maximum intensity level was a flux of about 1015 µmol/m^2^/s and for blue-shifted LEDs about 1400 µmol/m^2^/s, yielding a combined flux capacity of around 2415 µmol/m^2^/s. Other light parameters included the light level duration and frequency of cycling. We reasoned that because cells in more densely growing cultures would experience, on average, lower amounts of light (due to shelf-shading), it may be beneficial to ramp-up light intensities over time. Thus, we also included 2 parameters to configure the slope of light ramping on each LED type.

Another key environmental variable is pH. Spirulina cultures are typically grown in alkaline conditions. Our search space included a pH range from 8.0 to 10.0 with pH control bands as small as 0.2 units. Indoor spirulina cultures are typically grown at a temperature of about 35°C with sufficient air flow for mixing. We allowed for bioreactor growth regimens with two temperature levels, between 20°C to 37°C, with a specified duration and frequency. In defining all parameter bounds we considered that optima for biomass potency may not match the optima for biomass yield or volumetric productivity (similar to what we had observed for light intensity). Having defined these parameters and associated ranges for configuration (**Supplemental Table 1**), we set out to conduct ML-guided optimization in a complex, 17-dimentional space.

A key consideration for any ML optimization process is defining the reward function. To optimize volumetric productivity, we reasoned that ideal harvest times for some culture conditions may not necessarily fall on the last day of the experiment. Fast-growing cultures that reached peak productivity at earlier timepoints may be more optimal, provided that enough biomass is produced to justify the increased labor cost of more frequent run initiation and harvesting. Thus, we scored volumetric productivity, as measured by GFP fluorescence, at each sampling timepoint in association with a constant labor cost (**Methods, Equation 2**). Then we took the maximum score for each run (**Methods, Equation 3**). To better account for week-to-week fluctuations and to compare outcomes between run sets, we adjusted these run scores using the run set standards, and then calculated each run’s fold-improvement relative to the global standard mean (**Methods, Equation 4**). We refer to this productivity score (the reward) as “performance.”

To model the relationship between run parameter configurations and performance, we applied a Bayesian black-box approach. Bayesian, black-box ML is an automated approach to the joint optimization of a complex set of choices using reward function outcomes. The “black box” aspect allows us to optimize the function of bioreactor parameters without requiring that we know the function in closed form or its derivatives. This allows for unbiased, noise-corrupted (stochastic) observations of performance; the only requirement is that the function of bioreactor parameters can be evaluated at any point in the defined search space. With this approach, a prior belief is prescribed over all the possible run outcomes. This model is then refined as run sets are completed, using non-parametric, Gaussian Process (GP) algorithms. The posterior model represents updated beliefs about the bioreactor parameters and configurations most likely to produce high performance. The model then maximizes over this updated belief, using batch-wise optimization strategies to select the next set of experiments.

We explored the 17-dimensional configuration space by iteratively carrying out weekly bioreactor run sets with an average of 60 bioreactors on approximately a 90-hour growth interval (**Figure 3A**). Culture conditions for each bioreactor were separately programmed and controlled according to an assigned configuration, informed by the modeling results from previous runs. To monitor week-to-week variation, we also ran at least two reactors under standard conditions in every run set. Each day, we collected a culture sample from every bioreactor to monitor GFP accumulation. At the end of the run set, we scored each run’s performance. Then we fed these results back into the model, refining the model’s beliefs and using a batch-wise optimization strategy to select configurations for the next run set.

**Figure 3:**
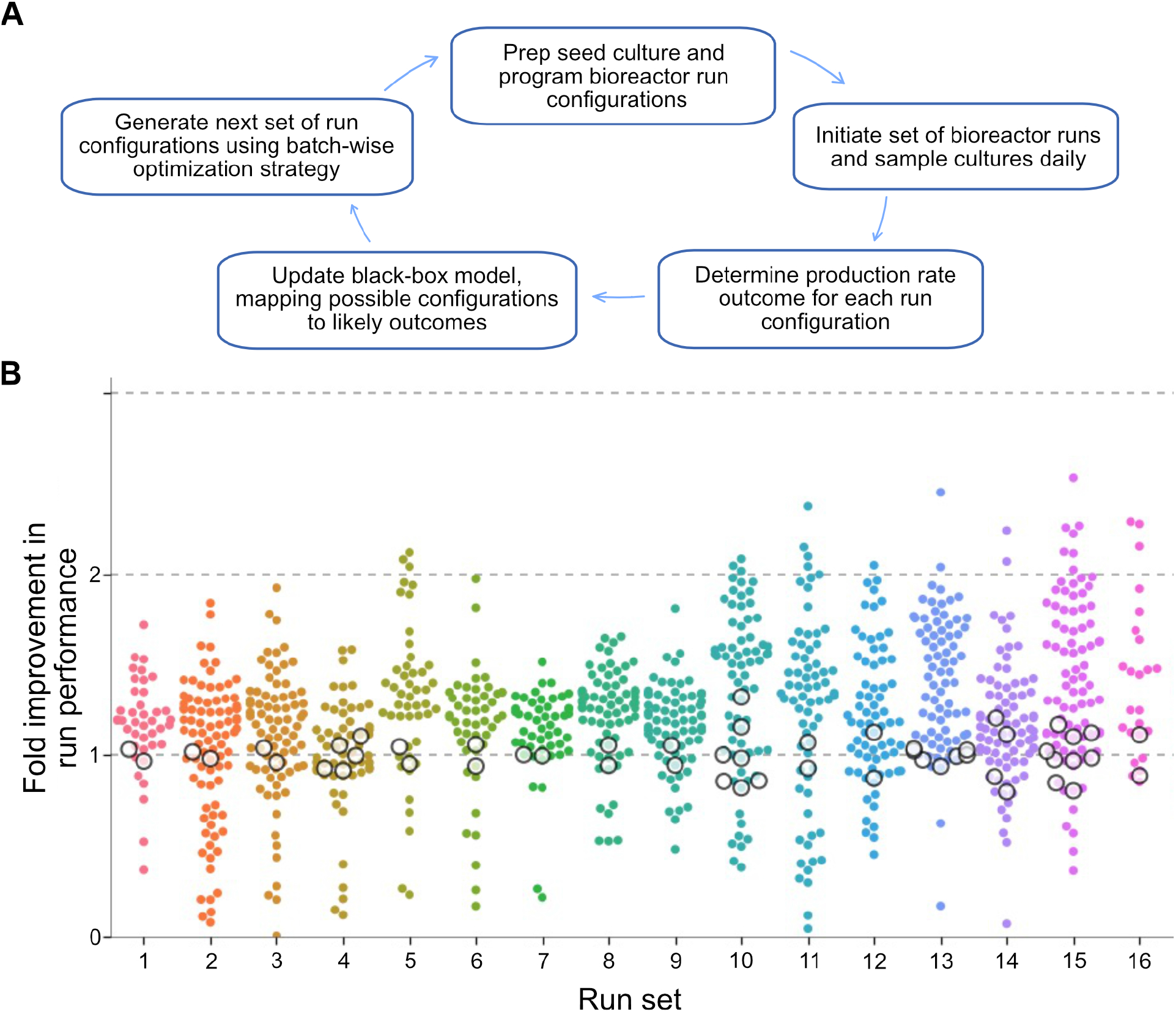
Bioreactor run performances over 16 rounds with targeted sampling of the run parameter search space. **A)** Workflow schematic for ML-guided optimization of bioreactor run performance. Each week ∼40-80 bioreactors are run with a set of parameter configurations for light, temperature, pH and mixing rate variables, totaling a 17-dimensional search space. **B)** Fold improvement in bioreactor run performance by round of ML-guided experimentation. Top plot shows individual bioreactor runs (dots) and standard condition runs (white circles) with fold improvement adjusted for run set variation in standards and calculated relative to the global mean of standards (1-fold, n = 51).

For early run sets, our batch-wise optimization strategy had a bias toward exploratory configurations. This allowed for broader sampling of the search space and broad mapping of the parameter configuration zones, helping identify potential performance peaks and valleys. We found that early run sets had a range of performance outcomes that suggested immediate improvements over standard runs. By run set number 5, we observed multiple run configurations with around a 2-fold performance improvement, despite having covered only a small proportion of the overall search space. Later, in run sets 11-16, we observed 15 additional configurations with greater than a 2-fold increase in performance (**Figure 3B**). To confirm the improvements, we took the initial top performers and ran a series of configuration replicates in run set 10. Improved configurations had consistently higher performance (group mean fold improvement = 1.8, standard deviation = 0.25, T test p-value = 3.3e-12) across 5 configuration replicates. Relative to standards, these replicates showed a stronger boost in signal after the first 24-hours of growth, leading to significant differences by 48-hours (T test p-value = 1.0e-06) (**Figure 4A top**). Mean volumetric productivity for each configuration was 69% to 93% higher than run set standards on the final day, leading to 81% higher volumetric yield on average (std = 25%) (**Figure 4A bottom**). For one of these high-performing configurations, we ran a second round of replicates in run set 13 and confirmed week-to-week robustness with a mean volumetric productivity improvement of 69% in the second run set (**Figure 4B**). Based on a conversion between GFP fluorescence units and µg GFP (**Methods, Equation 5**), we estimated the mean overall rate of protein production increased from 7.8 μg/mL/day to 14.2 μg/mL/day.

**Figure 4:**
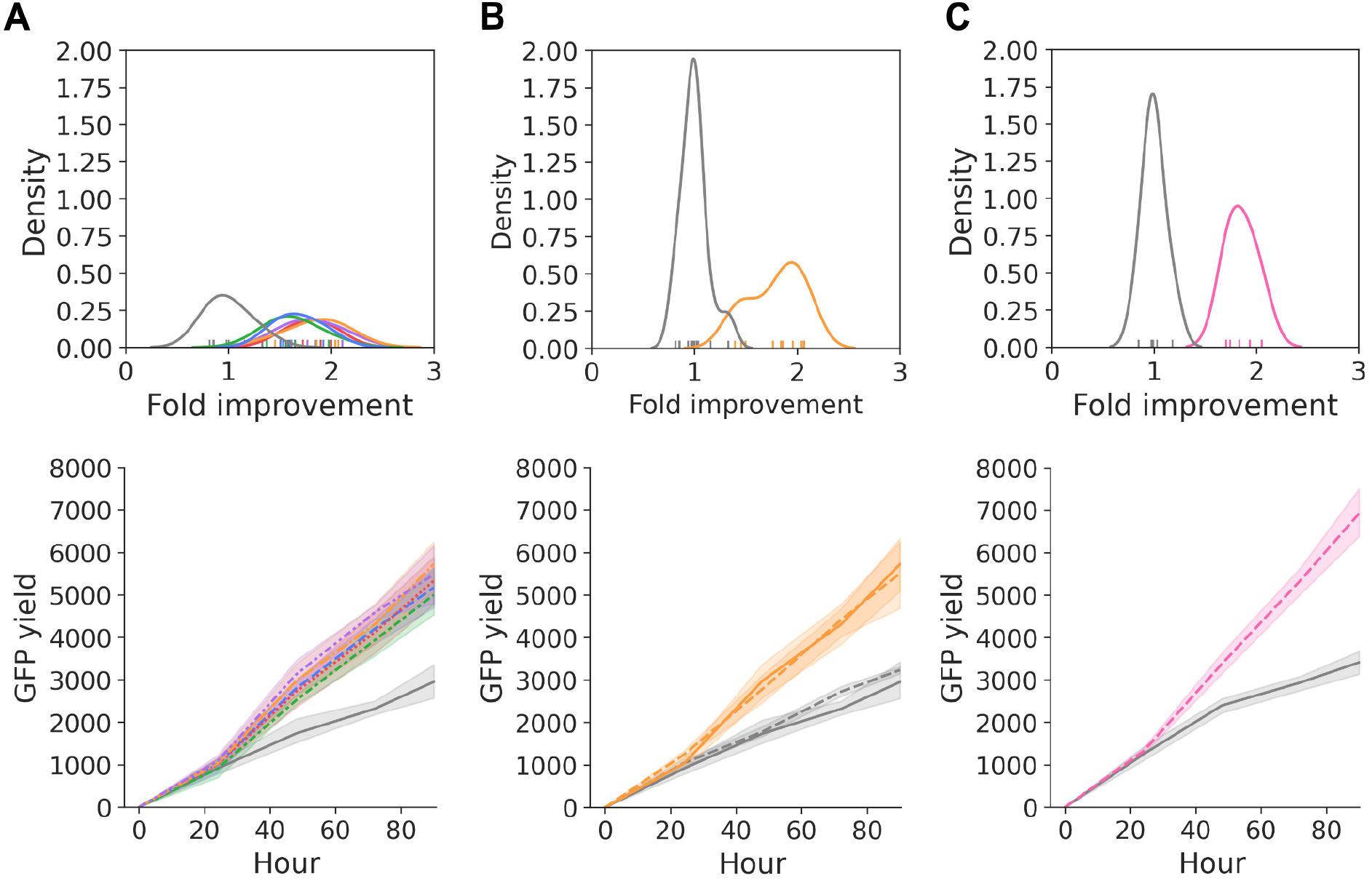
Kernel density estimate of fold-improvement (top) and time series of mean GFP fluorescence yield (bottom) for replicate runs of top-performing bioreactor configurations compared to standards (gray). **A)** Early, top-five run configurations in a validation run set (n = 5 each configuration, gray standard n = 7). **B)** Early top-performing configuration across two run sets (orange configuration n = 10, standard n = 14). **C)** Later-stage, top-performing configuration (pink configuration n = 5, blue standard n = 6). Shaded areas represent 95% confidence intervals.

For the second group of configurations in run sets 11-16, we took the top performer and sought to validate the discovery with 5 configuration replicates in run set 15. Once again, we confirmed that the high-scoring configuration had consistently better performance (mean fold improvement = 1.8, standard deviation = 0.14, T test p-value = 1.2e-06) (**Figure 4C top**). Mean volumetric productivity was 102% higher than the run set standards on the final day (std = 27%) (**Figure 4C bottom**). Thus, the model discovered multiple run configurations with performance outcomes that were more than 4 standard deviations away from our standard conditions, and these improvements were robust to replication, both within a given run set and across run sets. We concluded that modeling was successful in guiding the efficient discovery of high-performing bioreactor conditions for spirulina-based production of heterologous protein.

### Temperature, pH, and light adjustments contribute to improved performance

We observed that many high performing configurations had similar parameter settings. To further assess the degree to which these shared characteristics were representative of top-performing configurations, we binned each run based on performance and examined the binned histograms for each individual parameter and for derived variables (e.g., lowest temperature across temperature levels) (**Supplemental Figure S3**). We found that the best performing runs had strong temperature, pH, and initial light intensity biases (**Figure 5A**).

**Figure 5:**
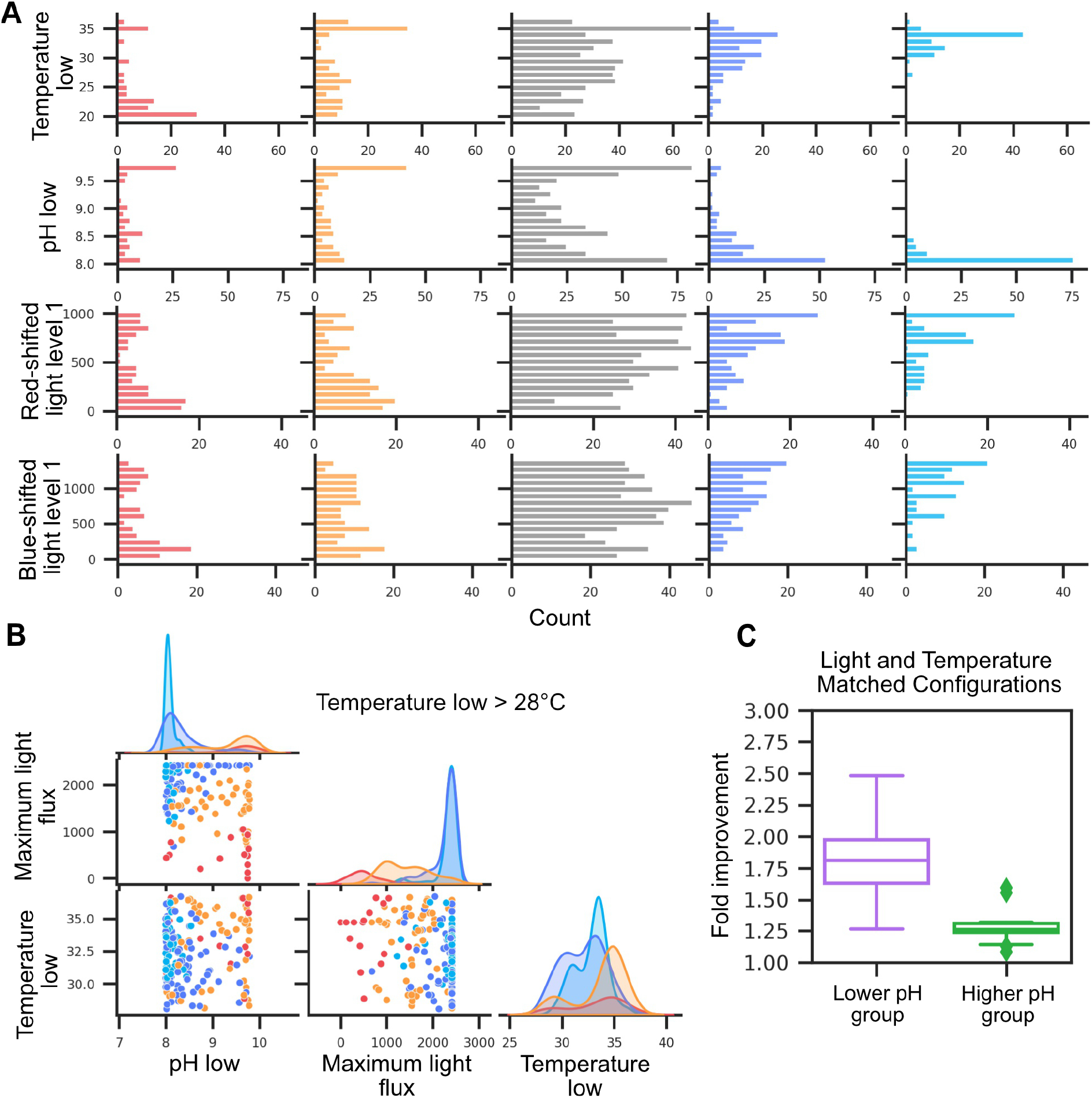
Highest-performing conditions tend to have similar pH, light, and temperature characteristics. **A)** Key single variable histograms binned by performance: lowest 10% (red), lower 25% (orange), within IQR (gray), upper 25% (dark blue), top 10% (teal). **B)** Key variable kernel density plots and paired scatter plots for run configurations with temperatures above 28°C. Run configurations are binned by performance: lowest 10% (red, temperature subset n = 16), lower 25% (orange, temperature subset n = 60), upper 25% (dark blue, temperature subset n = 100), top 10% (teal, temperature subset n = 66). **C)** Boxplot comparison of low pH configurations (purple, n = 65, pH lower bound < 8.55, pH upper bound < 9.23) and high pH configurations (green, n = 18, pH lower bound > 9.5) with temperature 33°C – 35°C and maximum light flux > 2000 µmol/m^2^/s.

Target temperatures for the top 10% of runs with constant temperature fell into a narrow band, with an interquartile range of 33.7°C to 34.1°C. The full range of sampled temperatures for runs in the top 10% spanned from 27.7°C to 36.6°C, suggesting that temperatures greater than ∼28°C may be necessary for the highest performance outcomes. Overall, the further away from a target temperature of 34°C, the stronger the association with lower run performance in our experiments. This “ideal” temperature fell below the maximum possible temperature of 37°C as well as the standard temperature of 35°C.

With respect to pH, the top 10% of runs displayed a strong bias for the lowest possible pH in our search space. These high-performing configurations had a median, lower bound pH of 8.06 and upper bound pH of 8.56 (lower IQR = 8.01 - 8.10; upper bound IQR = 8.31 - 8.63). Unlike temperature, where the highest-performing configurations had settings adjacent to our initial standard, the pH bias fell at the far end of the parameter search space from our initial standard pH of 9.75 to 9.95. While it was possible for a given run configuration to sample a wide-ranging pH, no runs in the top 10% exceeded an upper pH bound of 9.23, and all reached a lower bound of at least 8.54. These results suggest that dipping below a pH threshold of around 8.5 may be necessary for the highest performance outcomes.

Due to specification of light gradient parameters within our search space, there was an overall bias towards light schedules that would reach our equipment’s maximum light intensity. Across all sampled configurations, only 13% had a maximum light flux that fell below 1250 µmol/m^2^/s. Among the top 10% of runs (n = 72), only 12.5% (n = 9) configurations had a max light flux below 2000 µmol/m^2^/s. Nevertheless, among these high-performers, 4 configurations had a max light flux between 1200 and 1500 µmol/m^2^/s. This suggests that while maximizing light is a strong contributing factor to more optimal performance, it may not be necessary for top performance.

To examine the relationship between parameters, we began by plotting the values for each parameter pairing in bins based on run performance (**Supplemental Figure S4-S6**). We also created plots that applied third parameter cutoff values. Since there were no top performing runs with a temperature below 28°C, we applied this as one of our cutoffs. We observed that the low-performing configurations below this temperature had maximum light fluxes and lower-bound pH settings that spanned a range of values with no clear relationship (**Supplemental Figure S7**). Above this temperature, however, configurations in the top 25% tended to have either low pH or the maximum achievable light intensity; configurations in the top 10% tended to have both (**Figure 5B**). These observations suggest that light flux and pH depend on a temperature threshold but are otherwise independent parameters that combine to generate the highest performance outcomes. To further evaluate the contribution of lower pH to overall performance, we compared two groups of high-light, moderate temperature configurations: one with lower pH and the other with high pH. Median fold improvement in performance was 0.53-fold greater for the lower pH group (**Figure 5C**). We thus conclude that the ideal temperature band for cultivation falls around 33 to 34°C, and that for run configurations with temperatures above ∼28°C, both high light intensity (schedules with max light flux reaching at least 1200 µmol/m^2^/s) and low pH (lower bound < 8.5) contribute to dramatic performance improvements.

### Key parameter adjustments lead to improved yield with therapeutic strain and larger-scale cultures

We applied one of the top-performing run configurations (**Supplemental Table 2**) to culturing a spirulina strain that expresses a single-domain antibody (VHH) specific to *Campylobacter jejuni* flagellin in the 450 mL reactors. We compared growth (as measured by ash-free dry weight) and VHH levels (as a percentage of total soluble protein) between our initial standard and ML-discovered configuration. We found that cultures of this anti-campylobacter strain grew better than GFP strain cultures under our standard conditions. Despite this higher baseline, total biomass yield for the anti-campylobacter strain was 39% higher in the ML-discovered conditions after ∼90 hours of growth (one-tailed T test p-value = 0.04), corresponding to overall growth rate of approximately 0.68 g/L/day vs. 0.49 g/L/day in the standard condition (**Figure 6A**). VHH levels were unchanged in biomass samples from the ML-discovered condition (**Supplemental Figure S8**), indicating that performance improvements with the ML-discovered condition came primarily from improved growth.

**Figure 6:**
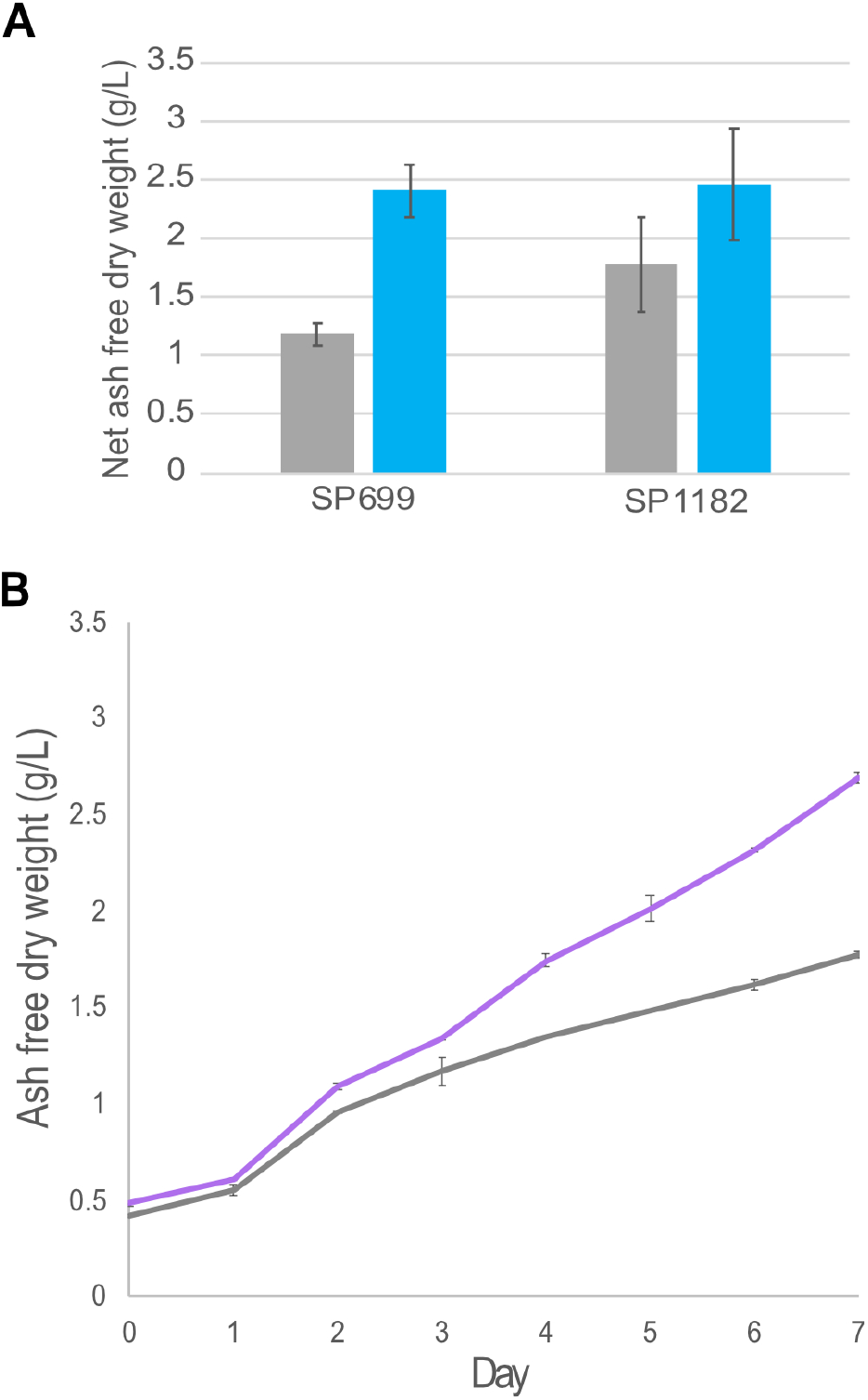
Application of high-performing conditions to anti-campylobacter strain in 450 mL and 250 L reactors increases biomass yields. **A)** Net biomass yield of the GFP-fusion (SP699) and anti-campylobacter VHH (SP1182) strains after ∼90 hours of growth at 450 mL scale. ML-discovered configuration (teal) compared with standard (gray). Error bars represent standard deviation of 3 run replicates. **B)** Biomass growth of an anti-campylobacter antibody strain (SP1182) in 250 L reactors. Improved condition based on ML-guided experimentation (orange) and initial standard condition (blue). Error bars represent standard deviation of AFDW measurements.

To confirm effect in a production-scale system, the anti-campylobacter strain (SP1182) was grown in parallel 250-liter flat panel photobioreactors under standard and improved conditions. We used a simplified, constant light program at 1350 µmol/m^2^/s, given constraints in the existing larger-scale system. We also had more approximate temperature controls than in the smaller-scale reactors. In a production run growth cycle totaling 7 days, the culture under improved conditions outperformed standard conditions, generating about 63% more biomass and higher VHH yields (**Figure 6B**). Thus, we conclude that lower pH (8.10 - 8.61) with higher light (1350 µmol/m^2^/s or more) and a slightly lower target temperature (33.3°C - 34.128°C) is a beneficial set of conditions to improve the yield of antibody-expressing strains at larger-scale.

## DISCUSSION

Traditional biologic platforms are ill-equipped to meet global demand, leaving a need for alternative, low-cost expression systems^2^. Indoor cultivation of spirulina provides an opportunity to finely tune growth conditions for low-cost production of biologics. We demonstrate that a Bayesian black-box model can efficiently steer cultivation conditions toward improved outcomes. Over iterative rounds of experimentation (between 5 to 15 rounds), we discovered multiple configurations with specifically tuned temperature, lower pH, and high overall light that approximately doubled productivities in a spirulina-based expression system. We further demonstrate that these conditions improve spirulina-based production of an anti-campylobacter antibody in large-scale culture.

Emerging biotechnology platforms introduce production organisms with unique and valuable attributes to the biopharmaceutical industry. A key challenge for emerging platforms is effectively evaluating multiple, often interacting environmental parameters that can impact productivity. Outside of ideal conditions, heterologous expression systems are particularly sensitive to stressors^21^. Thus, precise tuning of culture conditions can help to push the limits of cellular productivity while also ensuring more reliable production cultures and processes. Given limited resources, biologics manufacturers must try to both discover optimal combinations and finely tune key variables. Methods that reduce the total number of trials and evaluate variable interactions have improved upon the conventional one-variable at a time (OVAT) approach to variable optimization. Most notably, DOE approaches have saved time and money across multiple phases of drug development, leading to widespread adoption in the pharmaceutical and biotechnology industries.

DOE involves pre-determined study designs based on goals (e.g., screening, response surface mapping, optimization), number of factors to be investigated, and total number of experiments^9^. Often DOE is carried out in multiple phases. An initial screening phase applies fractional factorial designs to test combinations of variables at 2 or 3 levels^22,23^. After selecting key variables an optimization phase may follow, which tests additional center and axial points to help estimate linear and quadratic response curves and thus better “tune” each variable^24^. The specific design of a DOE-based study often takes a reasoned approach to characterizing complex, multivariate space by cautiously selecting appropriate variable levels and fractional factorials, applying both existing expert knowledge and thoughtful consideration.

In contrast with the pre-planned trials and separate phases for screening and optimization in many DOE studies, the adaptive nature of BO allows for greater flexibility in design. It is more robust to observation noise and variables that may interact non-linearly, such as genetic perturbations. BO samples broadly and efficiently across a landscape of possible variable combinations, continuously updates model assumptions based on the available data, and then adaptively shifts resources to conduct additional experiments in the most promising areas of a search space. With this adaptive modeling approach, researchers can rapidly move toward improved outcomes based on available information.

It can sometimes be difficult to appreciate the full value of an adaptive model in retrospect. As part of our study, we ran just 245 configurations prior to discovering multiple configurations that landed in the top 10% of performers overall. A comparable number of trials could be run in a 2-level, 8 factor, full factorial DOE study (2^8^ = 256 trials) by testing only a subset of variables, similar to our “simple” 8-parameter search sub-space (see **Methods**, “**Parameter space of controllable bioreactor alternatives**”). This traditional approach may have delivered similar directional information regarding temperature, pH, and light intensity responses, but reaching these conclusions would have depended on which variable levels were selected initially for testing. We found that temperatures for top performing bioreactor run configurations tended to fall in a narrow band around 33 to 34°C; DOE screening with a high temperature point of 37°C and a middle or low point between 25°C and 30°C, for example, would have missed this effect. Similarly, we found that all top-performing configurations had a pH range that fell below 8.5 units; DOE screening based on the spirulina literature, which widely reports using pH values between 9 and 10^25^, would likely have missed this pH effect if the low point was above 8.5. By de-risking the design phase selection of variable subsets and levels for testing, the adaptive model encourages broader exploration of possibilities, including variables that may contribute to higher-order interactions and non-convex responses. Furthermore, the rigidity of DOE would have presented significant limitations in sampling from both ramping and cyclic light schedules as a complex subspace, which in our study totaled 17-dimensions, requiring over 130,000 trials to complete a two-level full factorial experiment.

The adaptive, BO approach has a long history in fields outside of biotechnology^26–30^. It originated in the oil industry as a means of predicting more optimal oil-drilling locations and has become a *de facto* tool at leading machine-learning centers for model hyperparameter tuning^17,18,29,30^. Within biotechnology drug discovery fields, BO approaches have successfully guided chemical synthesis and protein engineering improvements^31,32^. Despite this history and demonstrated value, BO has had limited overall adoption in biotechnology, and it has not to our knowledge been applied in biologic manufacturing^33,34^.

We conclude that an adaptive, ML-based approach to optimization of culturing conditions is a valuable tool for biotechnology. BO can identify beneficial variable combinations and adjustments that might not otherwise be discoverable within high-dimensional data. As a complement to commonly applied DOE approaches, the adaptive ML approach can be especially helpful in exploring relatively uncharacterized systems, fine-tuning parameters in the context of many variables, and overcoming preconceived notions about an established system by using a wider search space with the potential for surprising discoveries. Future efforts could focus on new parameters, such as media and genetic variables, and could apply deep learning tools to incorporate other data sources, such as microscopy images, for further improvements in model prediction and search efficiency.

## ACKNOWLEDGEMENTS

This work was supported in part by funding from the Bill & Melinda Gates Foundation. We thank Craig Behnke, Troy Paddock, Mark Heinnickel, Lauren Goetsch, Thomas Adame, Ben Jester, Hui Zhao, Nhi Khuong, Zan Armstrong, Patrick Riley, Omar Vandal, and Dan Wattendorf for ongoing help, discussions, and advice. We thank Jake Siegel for review and editing.

## Supplemental Materials

### METHODS

#### Plasmids and strains

Strains were built using integration vectors directed toward a neutral integration site (Q01210 - Q01230 in the case of SP699) or D01030 *kmR* locus (in the case of SP1182). Integration vectors were built with a constitutively active, native promoter from upstream of spirulina’s *cpcB* gene (Pcpc600), as well as with the appropriate transgene, the *E. coli rrnB* terminator, a selection marker expression cassette, and 1-1.5 kb homology arms. SP699’s GFP transgene was constructed using Enhanced GFP (eGFP) fused to a synthetically designed, homo-dimeric scaffold and poly-histidine tag at the C-terminus (AA276). Methods for construction of SP1182 are further described in Jester, *et al*., 2020. Cultured cells were transformed into wild-type UTEX (SP003) or kanamycin resistance knock-out (SP205) and genotyped as described in Jester, *et al*., 2021. Pre-transformation cultures were grown in Multitron incubators at 35°C, 0.5% CO_2_, 110-150 μEi of light, and shaking at 120-270 rpm. Longer-term cultures were maintained in Innova incubators at 30°C, atmospheric C0_2_, 50-110 μEi of light, and shaking at 120 rpm. These small-scale cultures of 3-100 ml were grown in SOT media supplemented with 2.5-5 µg/ml streptomycin (for SP699) or 70-100 µg/ml of kanamycin (for SP1182) based on the transgene selection marker.

#### Bioreactor design (450 mL reactors)

Experiments were conducted in 96 independently controlled small, 450 mL, airlift reactors. All reactors were equipped with silicone adhesive-mount heater and a Neptune Systems temperature probe. A subset of the reactors were equipped for cooling with aluminum cooling heat sink plates. Each reactor was also equipped with a dimmable, dual color, LED-backlit LCD panel (Reefbright, NJ, USA) capable of illuminating the culture up to 2415 µmol/m^2^/s (3000 to 6500 Kelvin) as well as a pH probe to drive feedback control of culture pH via CO_2_ injection to the airlift stream. Each reactor is lit from the narrow ends to simulate a larger form factor flat plate bioreactor with commercially usable volume. Neptune Systems controllers with Apex software were used to monitor the temperature and pH and to control each reactor’s heating/cooling elements, solenoid for CO_2_ injection, and light panel intensity level.

#### Bioreactor run set experiments (450 mL reactors)

Seed train cultures used for inoculation were maintained in 9 L bioreactors in 1x SOT with 2x nitrate and sodium bi/carbonate buffer under the following settings: constant 500 µmol/m^2^/s, 34.8 - 35.3 °C, pH of 9.8 – 10, 2 LPM air flow for mixing. Three days prior to a run set, seed train cultures were harvested and used to re-inoculate seed reactors at a consistent inoculation density based on chlorophyll content.

To prepare inoculums, cells were transferred from seed reactors into an induction tank and allowed to settle. Liquid was decanted off the top, leaving a cell slurry. Inoculum slurries were assayed for chlorophyll content levels by first taking a 10 mL sample of slurry, centrifuging the sample for 10 min at 4,000 RPM, decanting, resuspending in DI water, transferring into 90 mL methanol, and placing in a sonicating water bath for 15 minutes. Then after taking a 1 mL aliquot of the methanol cell mixture and spinning it down, a 0.5 mL aliquot of supernatant was diluted and used to measure chlorophyll absorbance at 664 nm. This absorbance measure was used to estimate the chlorophyll content (12.1 μg/Abs unit) and biomass (total μg of chlorophyll / 1.8% of total biomass) and to calculate a volume of cell slurry for each bioreactor inoculation, targeting an initial estimated biomass of 500 mg/L.

Prior to bioreactor run initiation, all pH probes were checked for pH drift between readings. Probes were recalibrated if the cumulative drift exceeded 0.15 pH units. Benchtop reactors were filled with 320 mL of water followed by 40 mL 10x SOT with 2x nitrate, 40 mL 1 M sodium bi/carbonate pH 9, and 0.5 mL of 100g/L antifoam. Once filled, the CO_2_ tank value was opened and set to 30 PSI and flow rate on each CO_2_ rotamer was set to 0.1 LPM. Following inoculation, all reactors were brought up to a final volume of 450 mL and adjusted to the appropriate air flow rate. During the run, any water lost to evaporation was replaced by topping up reactor volumes each morning.

#### Bioreactor schedules and monitoring (450 mL reactors)

All bioreactor runs were registered in a PostgreSQL database together with a run configuration ID. Updated light and temperature settings were pushed to controllers every 12 minutes. Temperature and pH were actively monitored for deviations outside of programmed ranges and a record of light settings was updated with successful/failed pushes to the controller. Alarm notifications were triggered when temperatures reached 1.5 degrees above or below programmed values, as well as when pH readings reached 0.2 units above or below or when there were failed attempts to push light setting updates after a retry. Runs with either sustained alarms or recognized process deviations were flagged and manually reviewed. Upon review, runs with significant deviations, which may have impacted the integrity of results, were excluded from further analysis.

#### Cumulative pH calculation

Spirulina cells fix carbon during photosynthesis at a rate that correlates with biomass growth, and over time this increases the pH of a photobioreactor. When the upper bound pH setting is reached, this triggers the solenoid opening for the CO_2_ sparging, which lowers the pH back down to the lower bound of the pH range. The result is a sawtooth pattern, consisting of a gradual ascending interval to the upper bound and a steep return to the lower bound. To estimate biomass growth during bioreactor run experiments, we tracked the cumulative change in pH between solenoid firings and extrapolated changes during CO_2_ sparging times. Starting 30 minutes after run initiation, changes between pH sensor data points were cumulative added together. During CO_2_ sparging, the change in pH was estimated using a linear regression based on the previous 6 hours of data.

#### GFP fluorescence readings and normalization

For high-throughput protein quantification in bioreactor run experiments with SP699, daily samples of 1 mL were taken from each reactor and diluted (typically, 5 to 17-fold) in Thermo Scientific 8-well dishes with SOT to within the linear range of readings on a Spectramax M2 plate reader. Using SoftMax Pro 5.2 we ran a protocol script with shaking. Measurements to quantify GFP per unit of volume were taken with 488 nm excitation at 512 nm emission with a 495 nm cutoff filter. Additional control measurements of GFP were taken with 488 nm excitation at 540 nm emission with a 495 nm cutoff filter. Cell autofluorescence, composed primarily of chlorophyll and phycocyanin, was measured with 370 nm excitation at 660 nm emission with no cutoff filter. We adapted the instrument’s 96-well layout to 8-well plates by measuring at four central positions within each well of the 8-well plate and averaging the results. To account for position-to-position variation in the instrument’s readings, we first found each position’s median time zero fluorescence reading, and then adjusted readings to account for the relative difference between positional medians and a global median baseline.

#### Overview of batched Bayesian optimization

Bioreactors were run in parallel for many iterations to characterize the effects and co-dependence of bioreactor parameter settings on spirulina growth and protein product yield. By measuring cells grown in a large matrix of different environmental parameter settings, the best conditions for the highest protein yields can be realized; but both selection of the search space, for parameter settings over which to optimize, and the efficient prioritization of search space exploration is an open problem.

In this work, the settings for each bioreactor parameter were guided by the black-box Bayesian optimization (BO). This methodology provides an automated approach to the joint optimization of a complex set of choices, such as bioreactor parameter configurations. The “black box” aspect of the optimization refers to the fact that we do not need to know the closed form or the derivatives of f(bioreactor parameters) to optimize it and can allow for unbiased noise-corrupted (stochastic) observations of bioreactor run outcomes; the only requirement is that we are able to evaluate f(bioreactor parameters) at any point in the configuration space of interest.

Specifically, we used the following procedures for optimizing bioreactor run outcomes via BO:

1. Mapping spirulina growth condition alternatives to an explorable bioreactor parameter space
2. Defining a reward function to optimize over the parameter space
3. Selecting a batch (“set”) of trials (bioreactor runs) from the current space modeled via Gaussian Process Batched Upper Confidence Bound (GP-BUCB)
4. Updating Bayesian model from noisy observations of bioreactor run outcomes
5. Running the loop: repeat (selecting (#3) => updating (#4)) until “done” Each of these steps is further outlined in the following subsections.

#### Parameter space of controllable bioreactor alternatives

Each bioreactor run in ML-model experiments had an assigned run configuration with settings for 17 different parameters (**Supplemental Table 1**). The use of contextual bandits (see Golovin, *et al*.) allowed for the top-level selection of a “simple” light and temperature 8-dimensional subspace, while still allowing for identification of more complex light and temperature schedules in the full 17-dimensional space, if the complexity provided improved bioreactor run outcomes. This strategy also encouraged the model to sample equally from simple and complex search subspaces initially (i.e., both were tried in equal proportion when the model had a uniform prior, allowing fast discovery of simple, improved bioreactor run configurations if the simple parameter settings were on-par or better than complex parameter settings).

Light parameters made up 9 of the 17 parameters. Light level parameter settings were specified in units of µmol/m^2^/s, whereas each type of LED backlighting on the bioreactor LCD panels was controlled on a power scale of 1 to 100%. To convert between units of µmol/m^2^/s and the equipment’s power settings, specified intensities were normalized by the maximum achievable intensity: 1015 µmol /m2/s for the red-shifted LEDs; 1400 µmol /m/2 for the blue-shifted LEDs. Each LED-type (red-shifted and blue-shifted) had independent level and gradient settings. However, for configurations with two, alternating light levels, the timing of light level changes was synchronized between LED types. A light period was defined as one cycle of level 1 and level 2 settings. Cycle frequency was specified by the “number of light periods” parameter, which specified the total number of equal duration periods across 96 hours. The “light level 1 fraction” specified the proportion of each period at light level 1 versus light level 2. Light gradients were specified as slopes corresponding to the daily, percent of max increase in light intensity. Thus, a gradient slope of 0.5 corresponded to a 50% increase above specified light intensities over 24 hours. In application, these light level changes were pushed to controllers every 12 minutes. Gradients were applied at the start of a run, and as a result, light intensities often reached and maintained the maximum setting at some point during the run.

Beyond light parameters, there were 5 temperature, 1 air flow and 2 pH related parameters. Similar to light parameters, the “number of temperature periods” specified the total number of equal duration periods across 96 hours, and the “temperature level 1 fraction” specified the proportion of each period at temperature level 1 versus temperature level 2. The “air flow” parameter was manually set to the appropriate LPM on air flow meter. The pH parameters defined a band within which the pH was allowed to vary cyclically. The “pH lower level” (Φ_lower_) defined the lower bound of this allowed pH window while the “pH upper fraction” (*f*) defined the height of the window relative to the maximum achievable pH (Φ_max_) of 10.0. When a culture reached the upper extent of the allowed pH window, due to biomass growth, the pH was adjusted via CO_2_ bubbling back to the lower level. Due to physical hardware limitations, the allowed pH window had a minimum achievable width, referred to as Δ_min_. We then defined the upper extent of the allowed pH window (Φ_upper_) as a function of the following:

Constants:

Φ_max_ = 10.0 The maximum achievable/allowed pH for our system

Δ_min_ = 0.2 The minimum achievable pH window/gap size

Given:

Φ_lower_ The lower bound of the pH window (“pH lower level”)

*f* The fraction of the maximum achievable window size to use (“pH upper fraction”)

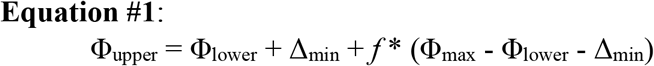

For example, given a pH lower level of 8.2 units, a pH upper fraction of 0.5 would place the upper-bound pH setting at 9.2 units; i.e., Φ_upper_ = 8.2 + 0.2 + 0.5*(10.0 - 0.2 - 8.2) = 9.2.

#### Reward function definition

The function being optimized in this case was: f(bioreactor parameters) = bioreactor run outcome; each reactor run having a unique configuration of parameter settings was equivalent to one evaluation of f(bioreactor parameters) and produced a resultant outcome at the end of each bioreactor run.

We defined a protein production curve for a single reactor over time as follows: Given:

F(*t*) The GFP fluorescence reading at time *t*

*C* Overhead cost of a production run

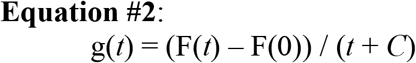

The cost parameter, *C*, encodes an overhead penalty associated with initiating and harvesting each production run cycle. We have used an empirically selected value of *C* = 200 in this work, as actual, live-production environment costs can carry complex sets of dependencies and uncertainties that are difficult to rapidly and accurately access. Given our protein production curve definition for a single run (Equation #2), we then defined the reward for a given bioreactor run as:

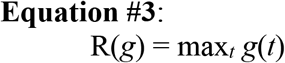

Each batch (“set”) of bioreactor runs was seeded by a common starting culture (see “Bioreactor run set experiments (450 mL reactors)”) and the interbatch performance variance was estimated by including multiple control condition replicates (“standards” based on initial conditions) within the batch to quantify both inter- and intra-batch variance. The reward outcome for a given bioreactor run was then adjusted to account for this inter-batch variance as follows:

Given:

µ_batch_ The mean intra-batch reward for our standard runs

µ_global_ The mean inter-batch (global) reward for our standard runs

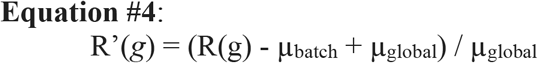

We then defined our per bioreactor run reward (termed “performance”) over which to optimize, f(bioreactor parameters), as this adjusted reward function: R’(*g*).

#### Selecting a batch of trials via GP-BUCB and iterating

The strategy employed in this work to search the defined parameter space was via Gaussian Process Batched Upper Confidence Bound (GP-BUCB) as implemented by Vizier^17^. The GP model (Bayesian black box model) prescribed a prior belief over all possible bioreactor run outcomes and its posterior represented updated beliefs about the bioreactor parameter settings most likely to produce the highest performance given the runs completed so far. Initially the model assigned a uniform prior belief across all outcomes. Selecting a single trial (bioreactor run) via the GP model can be performed by taking the bioreactor configuration (i.e., a point in model belief space) expected to produce the highest performance according to the current belief, as modeled by the GP according to the Upper Confidence Bound (UCB). However, extending this procedure to produce a batch of parameter choices optimally is an open problem.

In this work, the Batched Upper Confidence Bound (BUCB) approach selected subsequent points in the parameter space by simulating the model belief posterior, assuming a pessimistic performance for all points included in the batch so far, and then reapplying the UCB selection policy on this updated posterior. Each subsequent bioreactor configuration choice was made serially by the model using this simulated approach, resulting in a batch of bioreactor configuration choices to attempt in parallel.

Given a new batch (set) of bioreactor configuration evaluations (actual run performances), the belief posterior was updated with the full set of run performances to-date. Subsequent batches of runs were then sampled and evaluated cyclically by repeating this GP-BUCB policy.

#### Ash-free dry weight (AFDW) and chlorophyll content

Biomass samples were taken from each bioreactor and spun down in tabletop centrifuge for 10-15 minutes. Supernatant was removed. Biomass was transferred to glass tubes, washed with 10 mL of 200 mM NaCl, and spun down for another 10-15 minutes. After removing the supernatant, biomass was placed in a drying oven at 100°C for 24 to 72 hours. Once dry, biomass was removed, allowed to cool, and weighed on microbalance to obtain the pre-ash weight. Biomass material was then placed in an ashing furnace at 550°C for at least 16 hours followed by cooling. Samples were then weighed to obtain the post-ash weight with which to subtract from pre-ash weights.

To determine chlorophyll content of dried pellets and cell lysates, samples were diluted 1:9 in methanol and incubated on ice for 5-10 minutes. After spinning for 10 minutes to separate blue pellet, the absorbance of supernatant fraction was measured at 664 nm.

#### Immunoassay protein quantification

Cell lysates were prepared by a freeze-thaw and bead-beating protocol. 10 mL cell samples were collected, pelleted, and washed with 100 mM sodium chloride + 100 mM sodium phosphate, pH 6.0. Washed pellets were stored at -80°C overnight; then thawed in 100m mM sodium phosphate on ice and transferred to tube with glass beads. Samples were run ∼3 times in bead beater at 5000 for 30 seconds, saving the supernatant and washing beads with sodium phosphate buffer. After removing a subsample to determine chlorophyll content and the AFDW equivalent per uL of cell lysate, samples were transferred to 2x SDS buffer.

Cell lysate and purified proteins (from SP699) were separated by standard SDS-PAGE electrophoresis and transferred to a nitrocellulose membrane. Blots were probed with a monoclonal mouse anti-histidine tag antibody (GenScript), rinsed, and probed with rabbit anti-mouse horseradish peroxidase (HRP) secondary antibody. Chemiluminescent substrate was added, and the blot was imaged and then quantified using ImageJ. Total band intensity vs. loaded protein calibration curves were prepared for the purified proteins, and the concentration of epitope tagged protein in the lysate calculated from band intensity and this standard curve.

Protein levels for later run sets with SP1182 were measured by capillary electrophoresis immunoassay (CEIA). First, a Bradford assay was carried out to quantify total soluble protein in each run sample. Samples were then diluted and loaded with 0.12 µg of protein per capillary.

Purified protein controls were loaded in the range of 0.004 to 0.03 µg per capillary in generating a standard curve. Materials included: 12-230 kDa Jess/Wes Separation Module (ProteinSimple), mouse anti-histidine tag primary antibody (GenScript), and a rabbit anti-mouse HRP-conjugated secondary antibody (Protein Simple). Peak analysis was performed using the Protein Simple Compass software.

#### GFP unit conversion

From immunoassay quantification of GFP protein in an SP699 culture, grown under our standard conditions, we found there were ∼12321 µg of GFP per unit of chlorophyll absorbance.

Similarly, there were ∼1297 GFP fluorescence units per unit of chlorophyll absorbance. Thus, we defined a rough conversion factor (c) of 9.5e-3 µg GFP per unit of GFP fluorescence. Assuming these relationships held constant for a given bioreactor run timepoint with associated GFP fluorescence readings, the GFP yield in µg/mL (y(t)) was estimated as follows:

Given:

F(t) The GFP fluorescence reading at time t

c Conversion factor for GFP fluorescence to µg using sample chlorophyll content

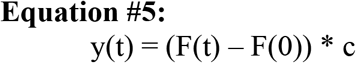

#### Bioreactor run experiments (250 L reactors)

To validate performance of BO improved conditions at scale, growth of the strains under standard and machine learning conditions was conducted in 250 L production reactors with similar light path and geometry. One reactor was run under standard conditions (pH 9.8 – 10.0; temperature 34.9-35.1 °C), the other reactor was run under conditions similar to high-performing run configurations from the BO search (pH 8.1-8.6, temperature 33.3-34.1 °C). Both reactors delivered the same light flux (1350 µmols/m²/sec) and the same media (1x SOT with 2x nitrate, identical to that described above in section “Bioreactor run set experiments (450 mL reactors)”. Cells were grown for seven days; relative growth rates are compared in Figure 6B.

## SUPPLEMENTAL TABLES

**Supplemental Table 1:**
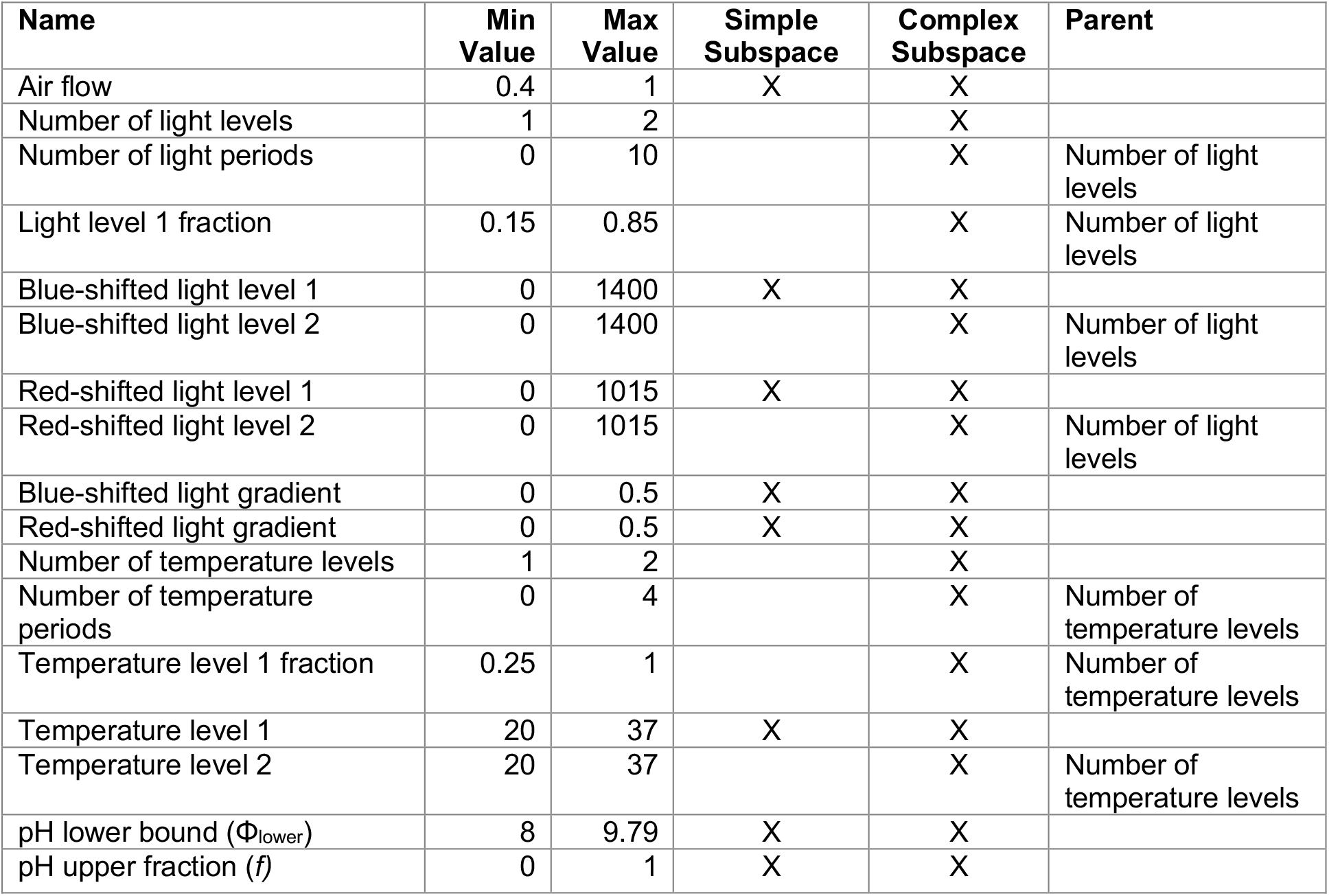
Model Parameters and Ranges

| Name | Min Value | Max Value | Simple Subspace | Complex Subspace | Parent |
| --- | --- | --- | --- | --- | --- |
| Air flow | 0.4 | 1 | X | X |  |
| Number of light levels | 1 | 2 |  | X |  |
| Number of light periods | 0 | 10 |  | X | Number of light levels |
| Light level 1 fraction | 0.15 | 0.85 |  | X | Number of light levels |
| Blue-shifted light level 1 | 0 | 1400 | X | X |  |
| Blue-shifted light level 2 | 0 | 1400 |  | X | Number of light levels |
| Red-shifted light level 1 | 0 | 1015 | X | X |  |
| Red-shifted light level 2 | 0 | 1015 |  | X | Number of light levels |
| Blue-shifted light gradient | 0 | 0.5 | X | X |  |
| Red-shifted light gradient | 0 | 0.5 | X | X |  |
| Number of temperature levels | 1 | 2 |  | X |  |
| Number of temperature periods | 0 | 4 |  | X | Number of temperature levels |
| Temperature level 1 fraction | 0.25 | 1 |  | X | Number of temperature levels |
| Temperature level 1 | 20 | 37 | X | X |  |
| Temperature level 2 | 20 | 37 |  | X | Number of temperature levels |
| pH lower bound ($\Phi_{\text{lower}}$) | 8 | 9.79 | X | X | |
| pH upper fraction ($f$) | 0 | 1 | X | X | |

**Supplemental Table 2:**
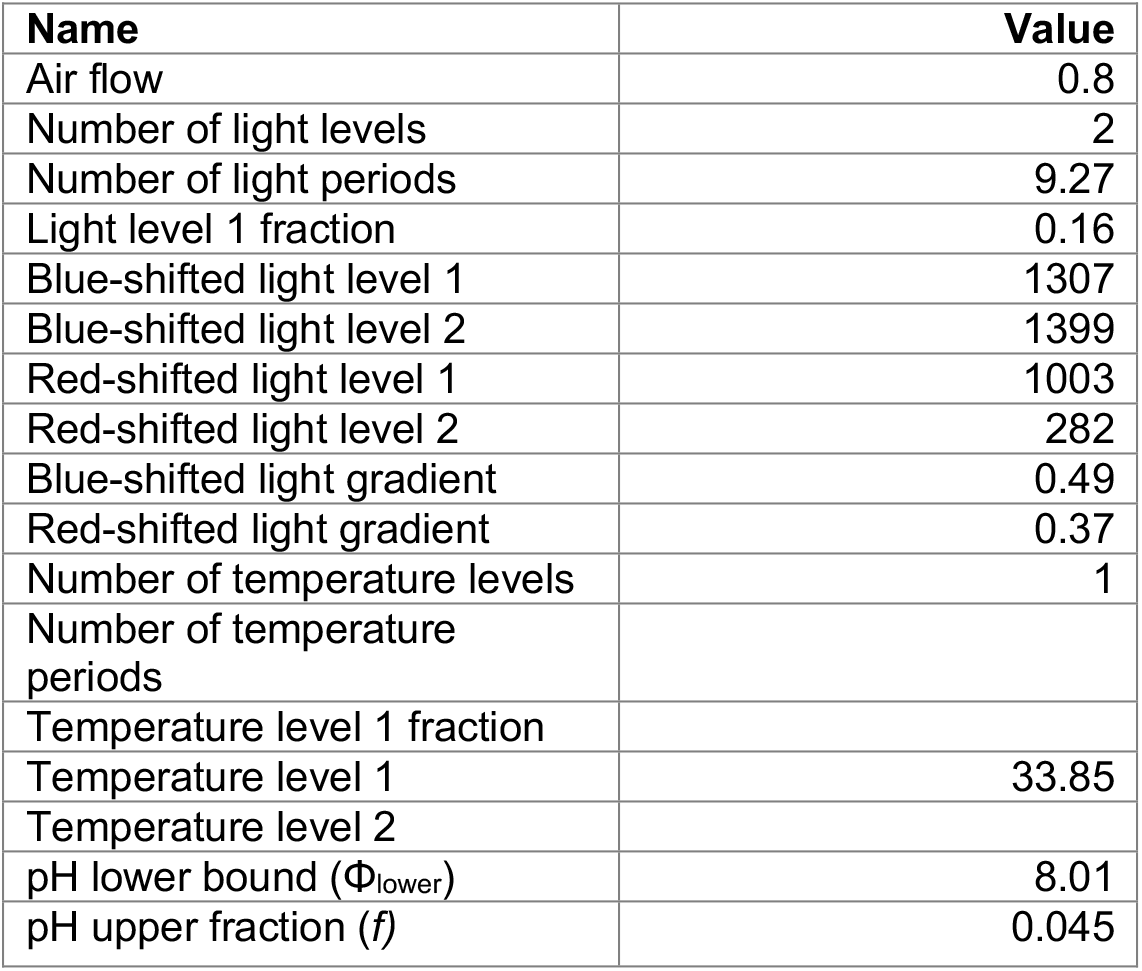
ML-discovered configuration applied to culturing a VHH-expressing strain, SP1182 (450 mL)

| **Name** | **Value** |
| --- | --- |
| Air flow | 0.8 |
| Number of light levels | 2 |
| Number of light periods | 9.27 |
| Light level 1 fraction | 0.16 |
| Blue-shifted light level 1 | 1307 |
| Blue-shifted light level 2 | 1399 |
| Red-shifted light level 1 | 1003 |
| Red-shifted light level 2 | 282 |
| Blue-shifted light gradient | 0.49 |
| Red-shifted light gradient | 0.37 |
| Number of temperature levels | 1 |
| Number of temperature periods |  |
| Temperature level 1 fraction |  |
| Temperature level 1 | 33.85 |
| Temperature level 2 |  |
| pH lower bound ($\Phi_{\text{lower}}$) | 8.01 |
| pH upper fraction ($f$) | 0.045 |

## SUPPLEMENTAL FIGURES

**Supplemental Figure S1:**
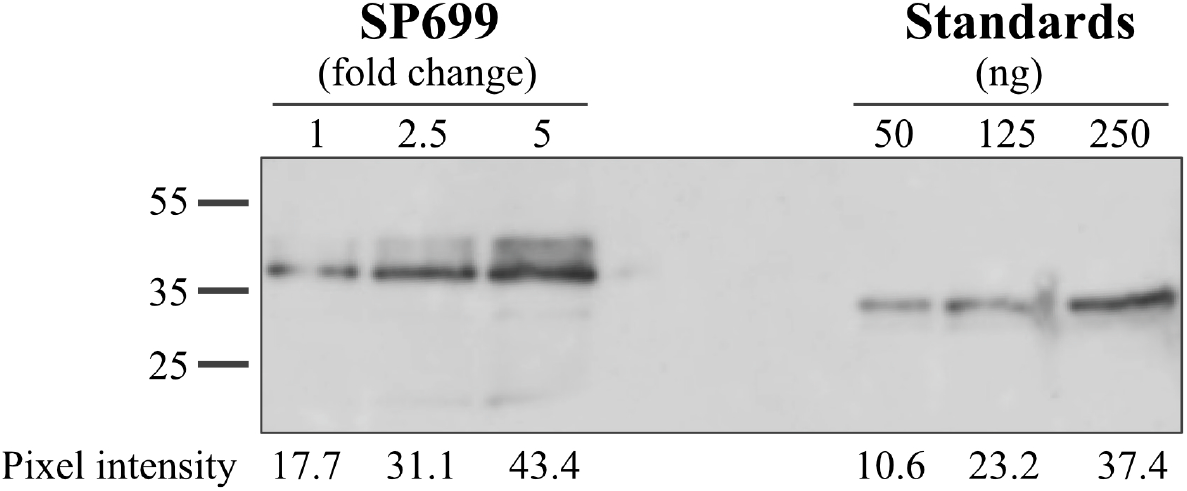
Western blot of his-tagged protein. Cell lysate from SP699 (GFP fusion strain) was loaded in three amounts and compared with the calibration curve of a known standard to quantify epitope-tagged protein in the lysate.

**Supplemental Figure S2:**
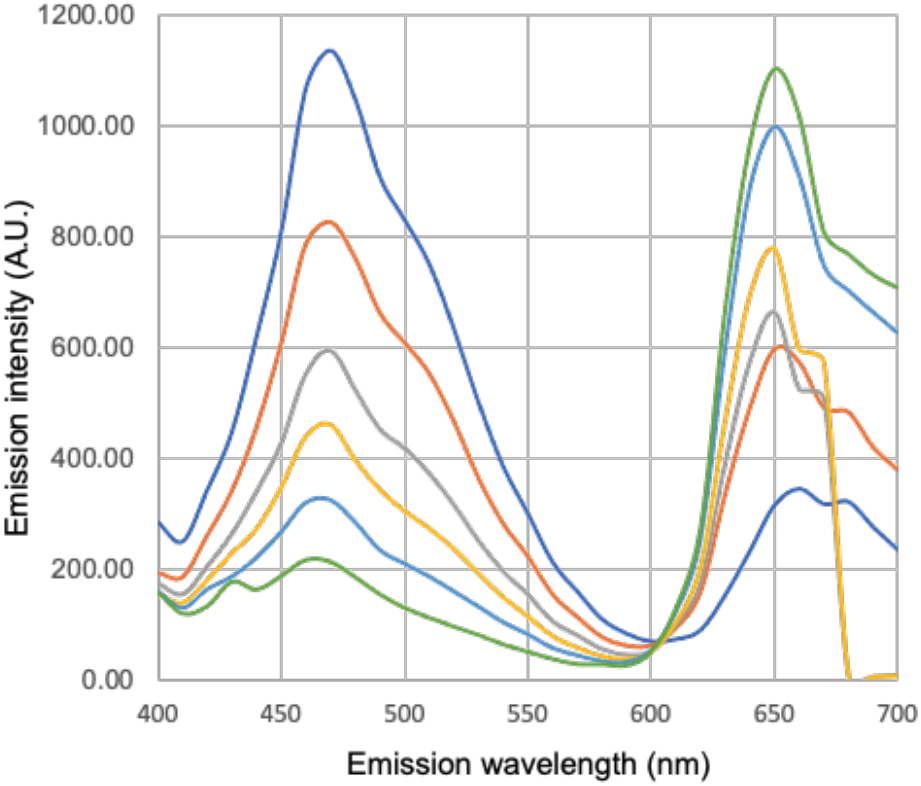
GFP and cell autofluorescence emissions have distinct peaks. The fluorescence emission spectra with 370 nm excitation shows one sample composed of 100% GFP-fusion (SP699) cells (dark blue) and samples mixed with WT (UTEX LB 1926) cells. Mixed samples are composed of 80% GFP-fusion (orange), 60% GFP-fusion (gray), 40% GFP-fusion (yellow), 20% GFP-fusion (light blue). The final sample is 100% WT (green).

**Supplemental Figure S3:**
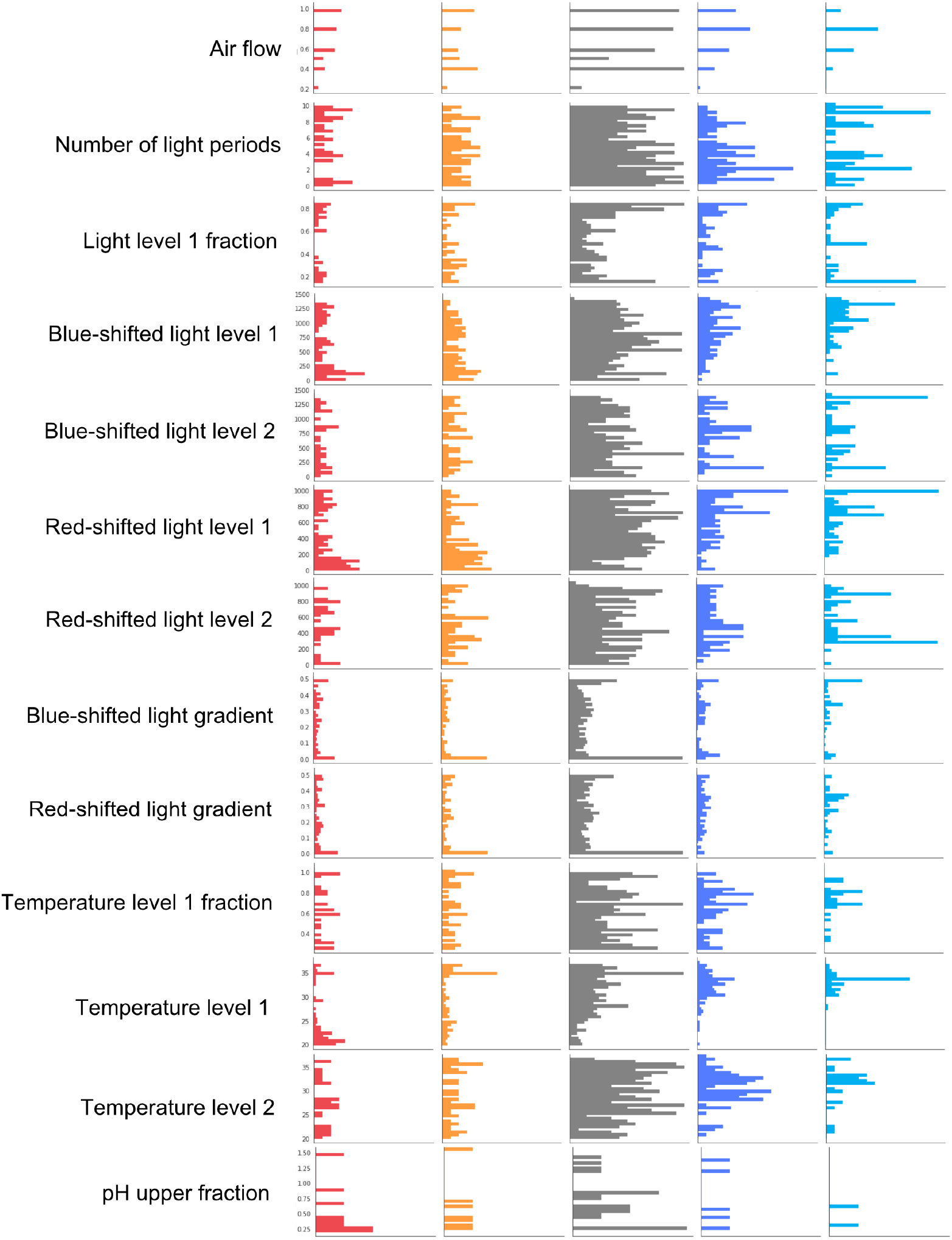
Subset of single parameter histograms by performance bin: lowest 10% (red), lower 25% (orange), within IQR (gray), upper 25% (dark blue), top 10% (teal).

**Supplemental Figure S4:**
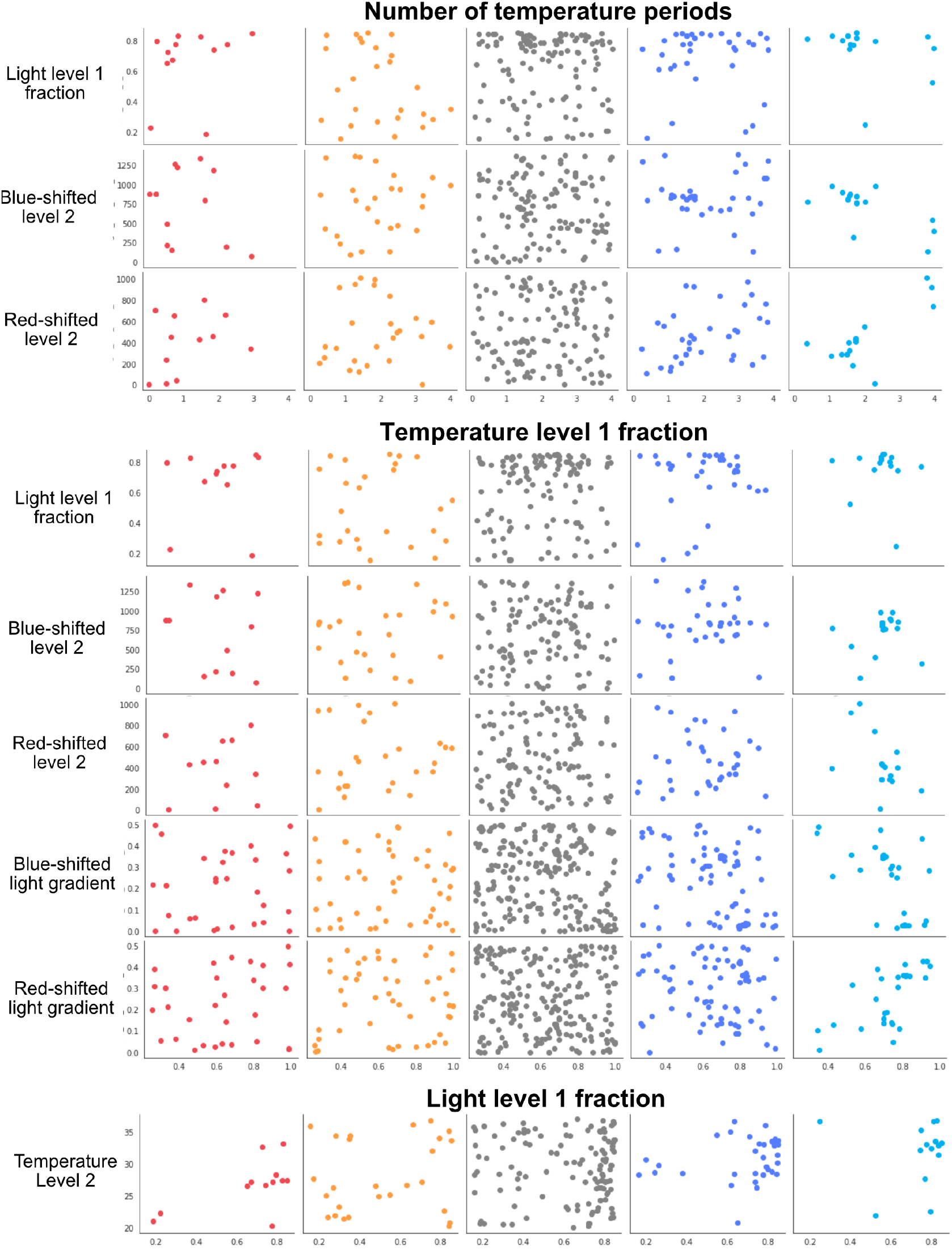
Selected subset of temperature-related parameter pairings by performance bin: lowest 10% (red), lower 25% (orange), within IQR (gray), upper 25% (dark blue), top 10% (teal).

**Supplemental Figure S5:**
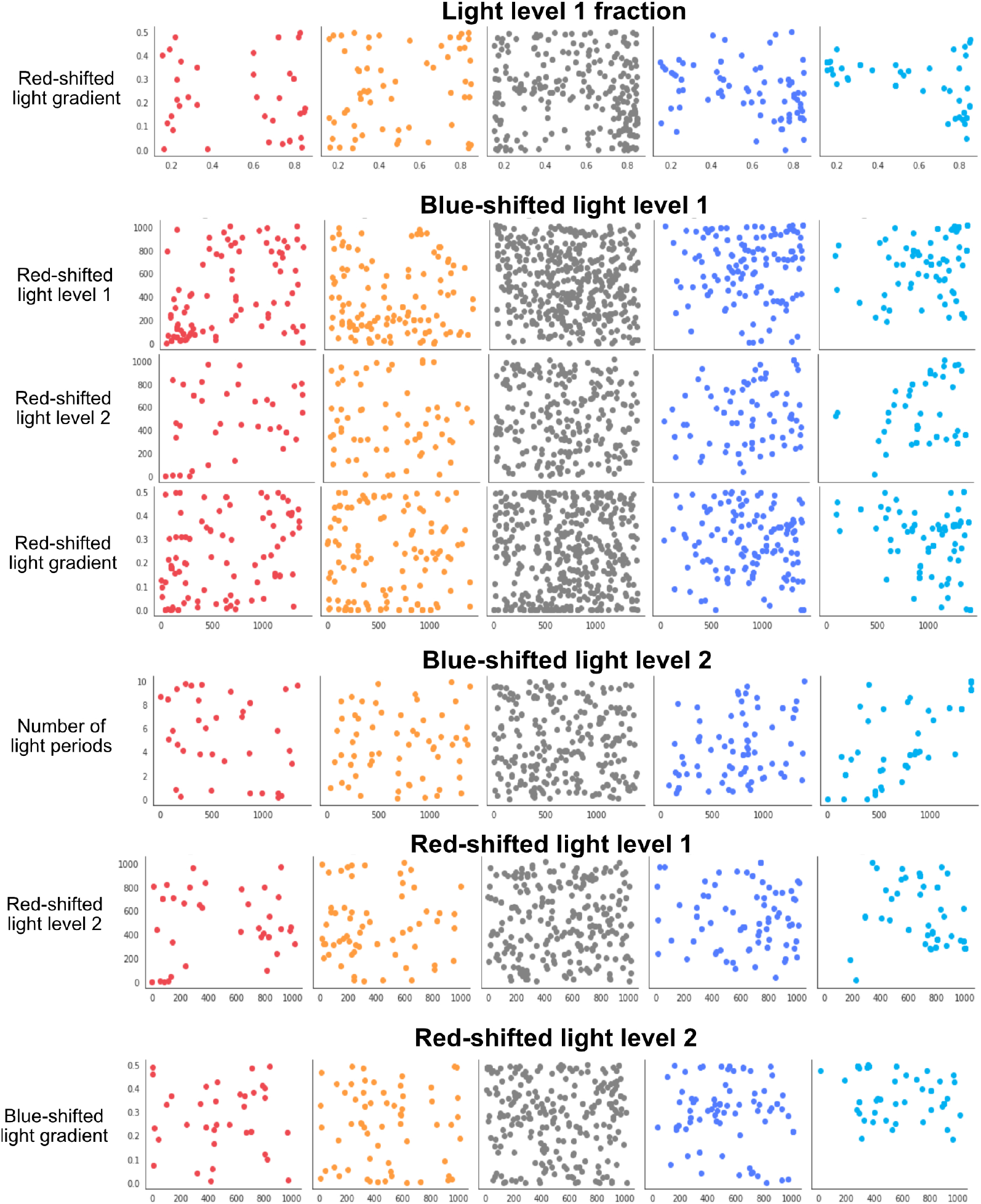
Selected subset of light-related parameter pairings by performance bin: lowest 10% (red), lower 25% (orange), within IQR (gray), upper 25% (dark blue), top 10% (teal).

**Supplemental Figure S6:**
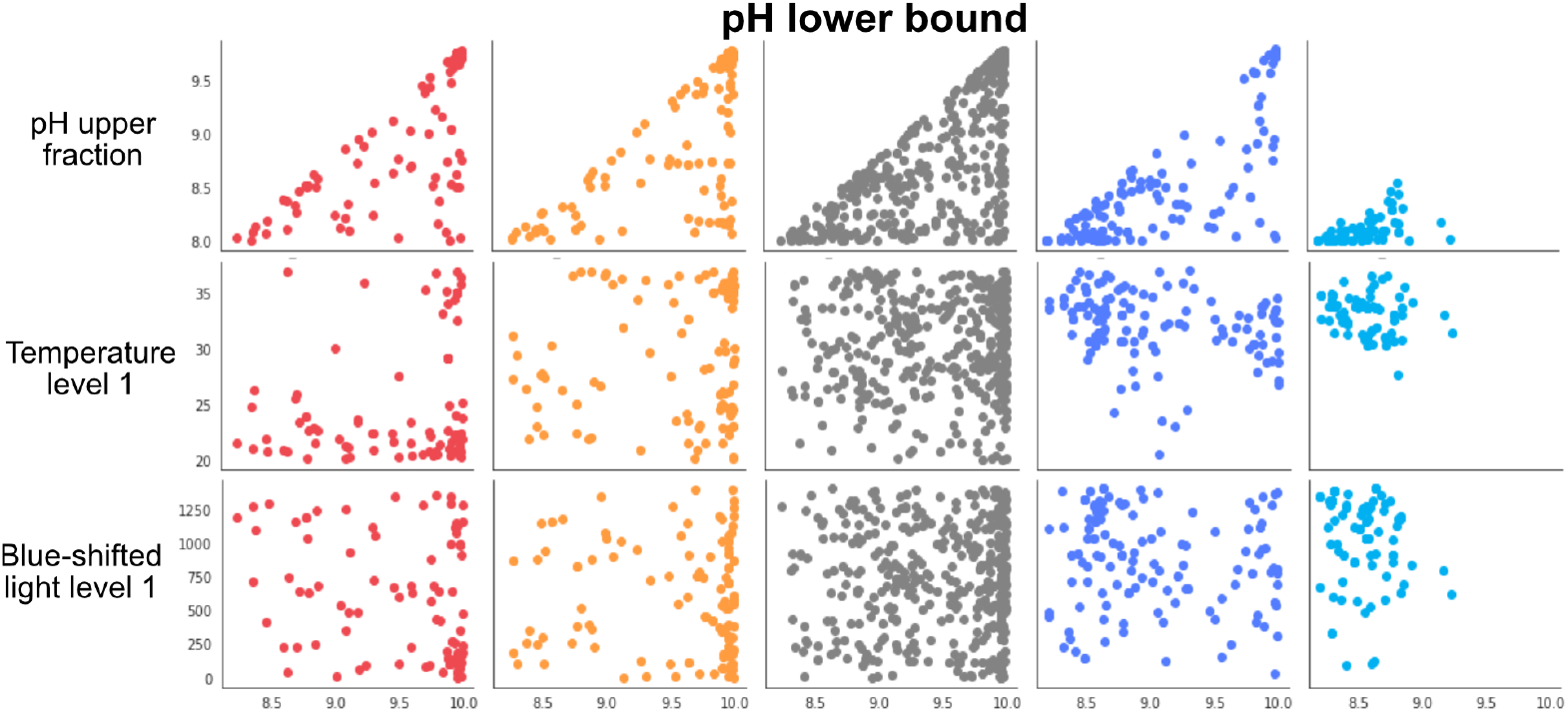
Selected subset of pH related parameter pairings by performance bin: lowest 10% (red), lower 25% (orange), within IQR (gray), upper 25% (dark blue), top 10% (teal).

**Supplemental Figure S7:**
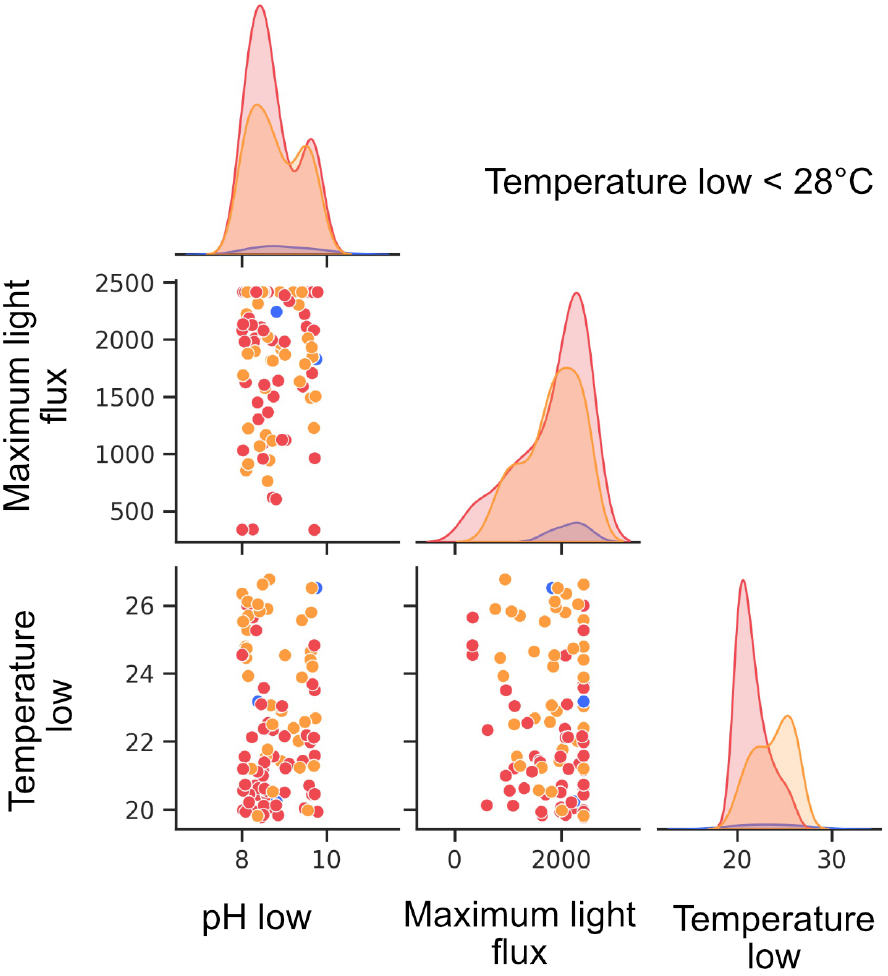
Key parameter kernel density plots and paired scatter plots for run configurations with temperatures below 28°C. Run configurations are binned by performance: lowest 10% (red, temperature subset n = 56), lower 25% (orange, temperature subset n = 42), upper 25% (dark blue, temperature subset n = 3). There are no configurations from the top 10% in this subset.

**Supplemental Figure S8:**
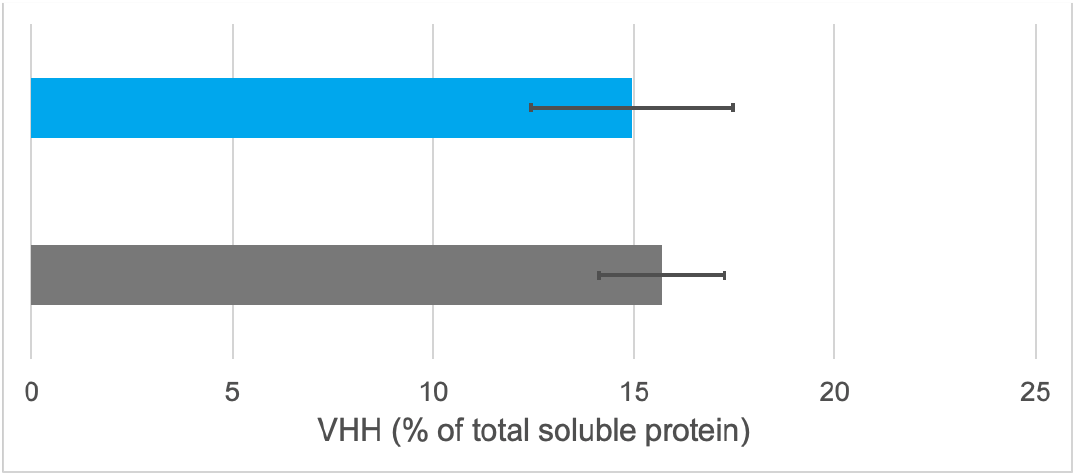
Mean VHH protein as a percentage of total soluble protein for biomass grown in ML-discovered configuration (teal) vs. standard (gray). Error bars represent standard deviation (n = 2 run replicates).

## REFERENCES

1. Merlin M, Gecchele E, Capaldi S, Pezzotti M, Avesani L. Comparative Evaluation of Recombinant Protein Production in Different Biofactories: The Green Perspective. BioMed Research International. 2014;2014:1–14. doi:10.1155/2014/136419

2. Ma JK-C, Drake PMW, Christou P. The production of recombinant pharmaceutical proteins in plants. Nat Rev Genet. 2003;4(10):794–805. doi:10.1038/nrg1177

3. Buyel JF, Twyman RM, Fischer R. Very-large-scale production of antibodies in plants: The biologization of manufacturing. Biotechnology Advances. 2017;35(4):458–465. doi:10.1016/j.biotechadv.2017.03.011

4. Vonshak A, ed. Spirulina Platensis (Arthrospira): Physiology, Cell-Biology and Biotechnology. CRC Press; 1997.

5. Gershwin ME, Belay A, eds. Spirulina in Human Nutrition and Health. 1st ed. CRC Press; 2007.

6. Marles RJ, Barrett ML, Barnes J, et al. United States pharmacopeia safety evaluation of spirulina. Crit Rev Food Sci Nutr. 2011;51(7):593–604. doi:10.1080/10408391003721719

7. Jester B, Zhao H, Gewe M, et al. Expression and manufacturing of protein therapeutics in spirulina. bioRxiv. Published online January 27, 2021:2021.01.25.427910. doi:10.1101/2021.01.25.427910

8. Kunert R, Reinhart D. Advances in recombinant antibody manufacturing. Appl Microbiol Biotechnol. 2016;100(8):3451–3461. doi:10.1007/s00253-016-7388-9

9. Singh V, Haque S, Niwas R, Srivastava A, Pasupuleti M, Tripathi CKM. Strategies for Fermentation Medium Optimization: An In-Depth Review. Front Microbiol. 2017;7:2087. doi:10.3389/fmicb.2016.02087

10. Powers DN, Trunfio N, Velugula-Yellela SR, Angart P, Faustino A, Agarabi C. Multivariate data analysis of growth medium trends affecting antibody glycosylation. Biotechnology Progress. 2020;36(1):e2903. doi:10.1002/btpr.2903

11. Lewis AM, Abu-Absi NR, Borys MC, Li ZJ. The use of ‘Omics technology to rationally improve industrial mammalian cell line performance. Biotechnology and Bioengineering. 2016;113(1):26–38. doi:10.1002/bit.25673

12. Xing Z, Li Z, Chow V, Lee SS. Identifying Inhibitory Threshold Values of Repressing Metabolites in CHO Cell Culture Using Multivariate Analysis Methods. Biotechnology Progress. 2008;24(3):675–683. doi:10.1021/bp070466m

13. Selvarasu S, Kim DY, Karimi IA, Lee D-Y. Combined data preprocessing and multivariate statistical analysis characterizes fed-batch culture of mouse hybridoma cells for rational medium design. Journal of Biotechnology. 2010;150(1):94–100. doi:10.1016/j.jbiotec.2010.07.016

14. Mora A, Nabiswa B, Duan Y, Zhang S, Carson G, Yoon S. Early integration of Design of Experiment (DOE) and multivariate statistics identifies feeding regimens suitable for CHO cell line development and screening. Cytotechnology. 2019;71(6):1137–1153. doi:10.1007/s10616-019-00350-1

15. Hill NR, Ayoubkhani D, McEwan P, et al. Predicting atrial fibrillation in primary care using machine learning. PLOS ONE. 2019;14(11):e0224582. doi:10.1371/journal.pone.0224582

16. Singal AG, Mukherjee A, Elmunzer JB, et al. Machine Learning Algorithms Outperform Conventional Regression Models in Predicting Development of Hepatocellular Carcinoma. Official journal of the American College of Gastroenterology | ACG. 2013;108(11):1723–1730. doi:10.1038/ajg.2013.332

17. Golovin D, Solnik B, Moitra S, Kochanski G, Karro J, Sculley D. Google Vizier: A Service for Black-Box Optimization. In: Proceedings of the 23rd ACM SIGKDD International Conference on Knowledge Discovery and Data Mining. ACM; 2017:1487–1495. doi:10.1145/3097983.3098043

18. Shahriari B, Swersky K, Wang Z, Adams RP, de Freitas N. Taking the Human Out of the Loop: A Review of Bayesian Optimization. Proceedings of the IEEE. 2016;104(1):148–175. doi:10.1109/JPROC.2015.2494218

19. Belay A. Mass Culture of Spirulina Outdoors–The Earthrise Farms Experience. In: Vonshak A, ed. Spirulina Platensis Arthrospira: Physiology, Cell-Biology And Biotechnology. CRC Press; 1997:131.

20. Wiltbank LB, Kehoe DM. Diverse light responses of cyanobacteria mediated by phytochrome superfamily photoreceptors. Nat Rev Microbiol. 2019;17(1):37–50. doi:10.1038/s41579-018-0110-4

21. Farkas Z, Kalapis D, Bódi Z, et al. Hsp70-associated chaperones have a critical role in buffering protein production costs. Wittkopp PJ, ed. eLife. 2018;7:e29845. doi:10.7554/eLife.29845

22. Grangeia HB, Silva C, Simões SP, Reis MS. Quality by design in pharmaceutical manufacturing: A systematic review of current status, challenges and future perspectives. European Journal of Pharmaceutics and Biopharmaceutics. 2020;147:19–37. doi:10.1016/j.ejpb.2019.12.007

23. Puente-Massaguer E, Badiella L, Gutiérrez-Granados S, Cervera L, Gòdia F. A statistical approach to improve compound screening in cell culture media. Engineering in Life Sciences. 2019;19(4):315–327. doi:10.1002/elsc.201800168

24. Al-Madboly LA, Khedr EG, Ali SM. Optimization of Reduced Glutathione Production by a Lactobacillus plantarum Isolate Using Plackett–Burman and Box–Behnken Designs. Front Microbiol. 2017;8. doi:10.3389/fmicb.2017.00772

25. Jung F, Jung CGH, Krüger-Genge A, Waldeck P, Küpper J-H. Factors influencing the growth of Spirulina platensis in closed photobioreactors under CO 2 – O 2 conversion. Journal of Cellular Biotechnology. 2019;5(2):125–134. doi:10.3233/JCB-199004

26. Li C, Rubín de Celis Leal D, Rana S, et al. Rapid Bayesian optimisation for synthesis of short polymer fiber materials. Sci Rep. 2017;7(1):5683. doi:10.1038/s41598-017-05723-0

27. Park S, Atwair M, Kim K, et al. Bayesian optimization of industrial-scale toluene diisocyanate liquid-phase jet reactor with 3-D computational fluid dynamics model. Journal of Industrial and Engineering Chemistry. 2021;98:327–339. doi:10.1016/j.jiec.2021.03.034

28. Lorenz R, Violante IR, Monti RP, Montana G, Hampshire A, Leech R. Dissociating frontoparietal brain networks with neuroadaptive Bayesian optimization. Nat Commun. 2018;9(1):1227. doi:10.1038/s41467-018-03657-3

29. Cressie N. The origins of kriging. Math Geol. 1990;22(3):239–252. doi:10.1007/BF00889887

30. Freund MJ. Cokriging: multivariable analysis in petroleum exploration. Computers & Geosciences. 1986;12(4):485–491. doi:10.1016/0098-3004(86)90063-4

31. Shields BJ, Stevens J, Li J, et al. Bayesian reaction optimization as a tool for chemical synthesis. Nature. 2021;590(7844):89–96. doi:10.1038/s41586-021-03213-y

32. Romero PA, Krause A, Arnold FH. Navigating the protein fitness landscape with Gaussian processes. PNAS. 2013;110(3):E193–E201. doi:10.1073/pnas.1215251110

33. Nikitin A, Fastovets I, Shadrin D, Pukalchik M, Oseledets I. Bayesian optimization for seed germination. Plant Methods. 2019;15:43. doi:10.1186/s13007-019-0422-z

34. Réda C, Kaufmann E, Delahaye-Duriez A. Machine learning applications in drug development. Computational and Structural Biotechnology Journal. 2020;18:241–252. doi:10.1016/j.csbj.2019.12.006

